# Common variants contribute to intrinsic human brain functional networks

**DOI:** 10.1101/2020.07.30.229914

**Authors:** Bingxin Zhao, Tengfei Li, Stephen M. Smith, Di Xiong, Xifeng Wang, Yue Yang, Tianyou Luo, Ziliang Zhu, Yue Shan, Nana Matoba, Quan Sun, Yuchen Yang, Mads E. Hauberg, Jaroslav Bendl, John F. Fullard, Panagiotis Roussos, Weili Lin, Yun Li, Jason L. Stein, Hongtu Zhu

**Author notes:** These authors contributed equally to this work (Bingxin Zhao, Tengfei Li). Corresponding author*: Hongtu Zhu, 3105C McGavran-Greenberg Hall, 135 Dauer Drive, Chapel Hill, NC 27599.

## Abstract

The human brain remains active in the absence of explicit tasks and forms networks of correlated activity. Resting-state functional magnetic resonance imaging (rsfMRI) measures brain activity at rest, which has been linked with both cognitive and clinical outcomes. The genetic variants influencing human brain function are largely unknown. Here we utilized rsfMRI from 44,190 individuals of multiple ancestries (37,339 in the UK Biobank) to discover and validate the common genetic variants influencing intrinsic brain activity. We identified hundreds of novel genetic loci associated with intrinsic functional signatures (*P* < 2.8 × 10^−11^), including associations to the central executive, default mode, and salience networks involved in the triple network model of psychopathology. A number of intrinsic brain activity associated loci colocalized with brain disorder GWAS (e.g., Alzheimer’s disease, Parkinson’s disease, schizophrenia) and cognition, such as 19q13.32, 17q21.31, and 2p16.1. Particularly, we detected a colocalization between one (rs429358) of the two variants in the *APOE* ε4 locus and function of the default mode, central executive, attention, and visual networks. Genetic correlation analysis demonstrated shared genetic influences between brain function and brain structure in the same regions. We also detected significant genetic correlations with 26 other complex traits, such as ADHD, major depressive disorder, schizophrenia, intelligence, education, sleep, subjective well-being, and neuroticism. Common variants associated with intrinsic brain activity were enriched within regulatory element in brain tissues.

---

The human brain is a complex system where functional organization and communication between brain networks are necessary for behavior and cognition^1–4^. The human brain remains active in the absence of explicit tasks or stimuli, resulting in an intrinsic functional architecture. Utilizing changes in blood oxygen level-dependent (BOLD) signal^5,6^, resting-state functional magnetic resonance imaging^7^ (rsfMRI) captures spontaneous intrinsic brain activity^8^. Specifically, the spontaneous neural activity and non-neural physiological processes within each functional region are quantified by the amplitude of low frequency fluctuations (ALFF) in BOLD time series^6,9,10^. Moreover, the inter-regional correlations in spontaneous neuronal variability are used to construct a functional connectivity matrix, which measures the magnitude of temporal synchrony between each pair of brain regions^6,11^.

rsfMRI has led to the discovery of multiple resting-state networks (RSNs) present in neurotypical human brains, including the default mode, central executive (i.e., frontoparietal), attention, limbic, salience, somatomotor, and visual networks^12–14^. Among these RSNs, the central executive, default mode, and salience networks are three core neurocognitive networks that support efficient cognition^15–17^. Accumulating evidence suggests that the functional organization and dynamic interaction of these three networks underlie a wide range of mental disorders, resulting in the triple network model of psychopathology^16,18^. Supporting this model, differences in RSNs have been detected in multiple neurological and psychiatric disorders^19^ relative to neurotypical controls, such as Alzheimer’s disease^20^, Parkinson’s disease^21^, and major depressive disorder (MDD)^22^.

Twin and family studies have largely reported a low to moderate degree of genetic contributions to intrinsic brain activity^23–29^. For example, the family-based heritability estimates of major RSNs ranged from 20% to 40% in the Human Connectome Project (HCP)^30^. In a previous study using about 8,000 UK Biobank (UKB) individuals^31^, the SNP heritability^32^ of amplitude and functional connectivity traits can be higher than 30%. Although there were multiple candidate gene studies for intrinsic brain activity (such as for *APOE*^33^ and *KIBRA*^34^), currently only one genome-wide association study (GWAS)^31^ has been successfully performed on rsfMRI^23^ (*n* ≈ 8000). This is likely due to both insufficient sample size for GWAS discovery and weaker genetic effects on brain function than structure^31,35–39^. It is also known that functional connectivity traits in rsfMRI are typically noisier than brain structural traits measured in other neuroimaging modalities. In addition, imaging batch effects^40^ (e.g., image acquisition, processing procedures, and software) may cause additional technical variability in rsfMRI analyses^41^, making GWAS meta-analysis and independent replication particularly challenging. Therefore, genetic variants influencing intrinsic brain activity have remained largely undiscovered and their shared genetic influences with other complex traits and clinical outcomes are unknown.

To address these challenges, here we collected individual-level rsfMRI data from four independent studies, including the UK Biobank^42^, Adolescent Brain Cognitive Development (ABCD^43^), Philadelphia Neurodevelopmental Cohort (PNC^44^), and HCP^45^. We harmonized rsfMRI processing procedures by following the unified UKB brain imaging pipeline^9,46^. Functional brain regions and corresponding functional connectivity were characterized via spatial Independent Component Analysis (ICA)^47,48^ for 44,190 individuals from multiple ancestries, including 37,339 from UK Biobank. As in previous studies^9,31,49^, two parcellations with different dimensionalities^13,50^ (25 and 100 regions, respectively) were separately applied in spatial ICA and we focused on the 76 (21 and 55, respectively) regions that had been previously confirmed to be non-artifactual^9^. Two group of neuroimaging phenotypes were then generated: the first group contains 76 (node) amplitude traits reflecting the regional spontaneous neuronal activity; and the second group includes 1,695 (i.e., 21 × 20/2 + 55 × 54/2) (edge) functional connectivity traits that quantify the inter-regional co-activity, as well as 6 global functional connectivity measures summarizing all of the 1,695 pairwise functional connectivity traits^31^. These 1,777 traits were then used to explore the genetic architecture of intrinsic brain activity. To aid interpretation of GWAS results, the functional brain regions characterized in ICA were labelled by using the automated anatomical labeling (AAL) atlas^51^ and were mapped onto major functional networks defined in Yeo, et al. ^14^ and Finn, et al. ^12^. Our GWAS results can be easily explored and downloaded through the Brain Imaging Genetics Knowledge Portal (BIG-KP) https://bigkp.org.

## RESULTS

### Genetics of the intrinsic brain functional architecture

SNP heritability was estimated for the 1,777 intrinsic brain activity traits via GCTA^52^. The mean heritability (*h^2^*) estimate was 27.2% (range = (10%, 36.5%), standard error = 6.0%) for the 76 amplitude traits, all of which remained significant after adjusting for multiple comparisons by using the Benjamini-Hochberg procedure to control false discovery rate (FDR) at 0.05 level (1,777 tests, **Fig. 1a** and **Supplementary Table 1**). Among the 1,701 functional connectivity traits, 1,230 had significant (again at 5% FDR) heritability with estimates varying from 3% to 61% (mean = 9.6%, standard error = 5.8%). Ten functional connectivity traits had heritability higher than 30%, including 4 global functional connectivity measures (**Supplementary Fig. 1**) and 6 pairwise functional connectivity traits (**Fig. 1b**). These most heritable traits were most related to the central executive, default mode, and salience networks in the triple network model of psychopathology^16^. To examine whether intrinsic brain activity within the triple network in general had higher heritability, we classified the 76 amplitude traits into two categories 1) fully or partially within the triple network and 2) outside the triple network. Correspondingly, the 1,695 pairwise functional connectivity traits were classified into 1) within the triple network, 2) outside the triple network, and 3) between the triple and non-triple networks. We found that amplitude traits within the triple network had significantly higher heritability than those outside the triple network (mean = 30.5% vs. 22.3%, *P* = 6.3 × 10^−11^, two-sided Wilcoxon rank test) (**Fig. 1c**). Similarly, functional connectivity traits within the triple network had higher heritability than interactions outside the triple network or between the triple and non-triple networks (mean = 12.5% vs. 7%, *P* = 1.9 × 10^−26^). These results indicate that the level of genetic control might be higher in core neurocognitive networks. The range of heritability estimates was consistent with previous results^31^, suggesting that common genetic variants had a low to moderate degree of contributions to inter-individual variability of intrinsic brain activity. The overall genetic effects on both amplitude and functional connectivity were lower than those on brain structure. For example, the average heritability was reported to be 48.7% for diffusion tensor imaging (DTI) traits of brain structural connectivity in white matter tracts^53^ and 40% for regional brain volumes measuring brain morphometry^37^. Nevertheless, as shown below, intrinsic brain activity may be more functionally relevant with stronger genetic connections to brain disorders than brain structure, such as Alzheimer’s disease.

**Figure 1:**
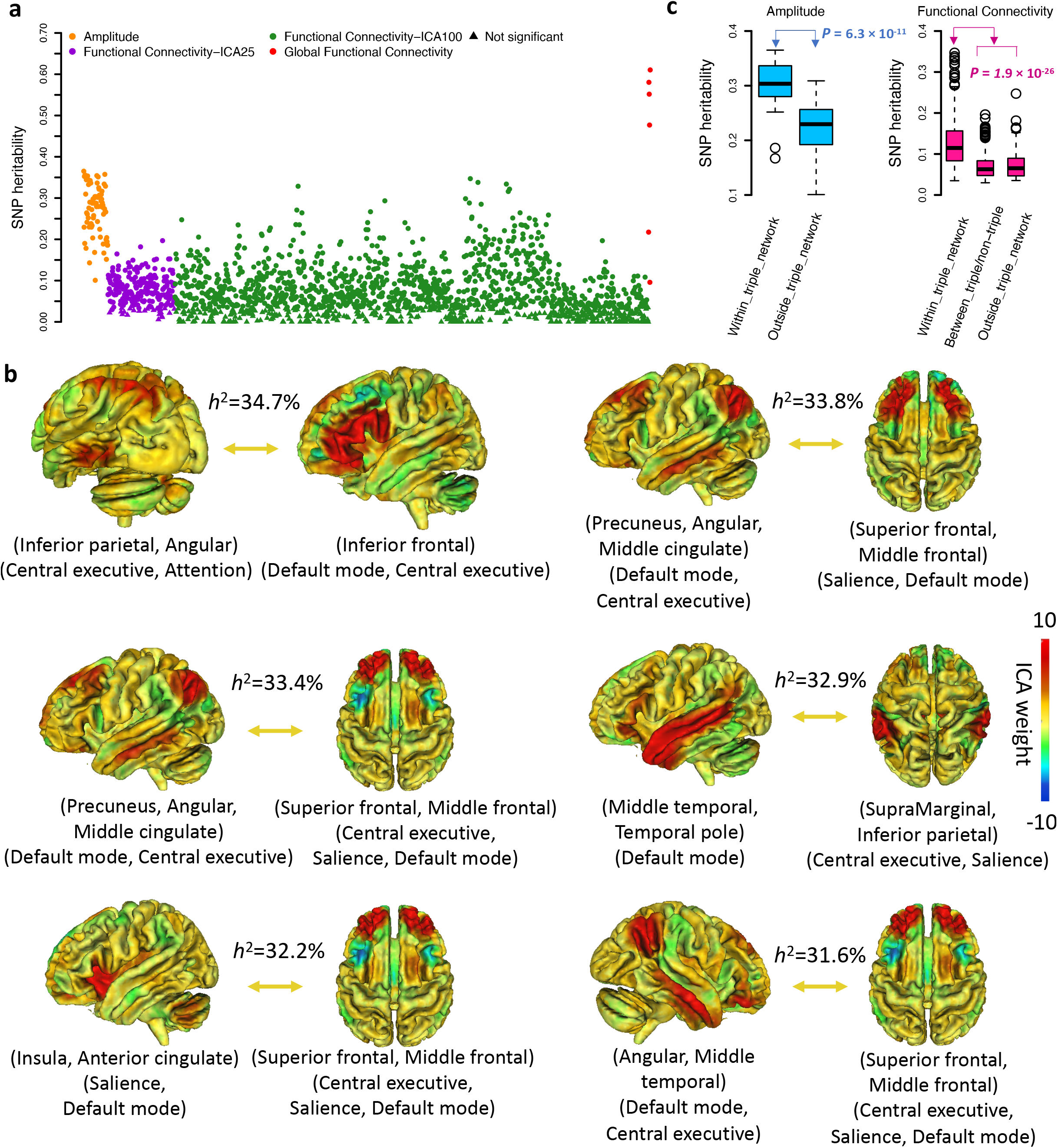
SNP heritability analysis of rsfMRI traits (n = 34,691 subjects). **a)** Heritability estimates of 1,777 rsfMRI traits of brain activity, including 76 amplitude traits, 1,695 pairwise functional connectivity traits (from two parcellations with 25 and 100 dimensionalities, respectively), and 6 global functional connectivity measures. **b)** Location and functional network of the pairs of functional regions (i.e., nodes) characterized by spatial independent component analysis (ICA) whose inter-regional functional connectivity had heritability (*h*^2^) higher than 30%. The color represents the weight profile of the ICA node. For example, the functional connectivity between two ICA nodes mainly over the inferior parietal, angular and inferior frontal regions had *h*^2^ = 34.7%. **c)** Comparison of the heritability within the triple network (i.e., the three core neurocognitive networks: central executive, default mode, and salience) and the heritability outside the triple network. *P*-value (*P*) of the two-sided Wilcoxon rank test was used to evaluate the difference.

Genome-wide association discovery was carried out for 1,777 intrinsic brain activity traits using UKB individuals of British ancestry (*n* = 34,691, Methods). The Manhattan and QQ plots can be found in the BIG-KP server. At the significance level 2.8 × 10^−11^ (5 × 10^−8^/1,777, i.e., the standard GWAS threshold, Bonferroni-adjusted for the 1,777 traits), FUMA^54^ identified 264 lead independent variants (linkage disequilibrium [LD] *r^2^* < 0.1), and then characterized 606 significant locus-trait associations for 197 traits (75 amplitude and 122 functional connectivity (**Supplementary Tables 2-3**, **Supplementary Fig. 2**, Methods). The amplitude traits typically had multiple associated variants and a number of variants were widely related to the amplitude in different brain regions, such as rs429358 (nearest gene *APOE*), rs2274224 (*PLCE1*), and rs1133400 (*INPP5A*). In addition, rs2279829 (*ZIC4*), rs62158211 (*AC016745.1*), and rs115877304 (*NR2F1-AS1*) were associated with multiple functional connectivity traits. Global and pairwise functional connectivity traits that had at least 5 significant variants were again most related to the central executive, default mode, and salience networks (**Supplementary Fig. 3**). Of the 14 associated variants that had been identified in the previous GWAS^31^, 12 were in LD (*r*^2^ ≥ 0.6) with our significant variants, most of which were associated with amplitude traits. In summary, our analyses identify many novel variants associated with intrinsic functional signatures and illustrate the global genetic influences on functional connectivity across the whole brain. The degree of genetic control is higher in the central executive, default mode, and salience networks, whose cross-network interactions closely control multiple cognitive functions and affect major brain disorders^18^.

### Replication and the effect of ancestry

We aimed to replicate our results in UKB British GWAS using other independent datasets. First, we repeated GWAS on UKB individuals of White but Non-British ancestry (UKBW, *n* = 1,970) and three non-UKB European-ancestry cohorts, including ABCD European (ABCDE, *n* = 3,821), HCP (*n* = 495), and PNC (*n* = 510). We meta-analyzed the four European GWAS (total *n* = 6,796) and checked whether the locus-trait associations detected in UKB British GWAS can be replicated. For the 606 significant associations, 101 (16.7%) passed the 8.2 × 10^−5^ (i.e., 0.05/606) Bonferroni significance level in this validation GWAS, and 599 (98.8%) were significant at FDR 5% level. Next, we performed GWAS on four non-European validation datasets: the UKB Asian (UKBA, *n* = 446), UKB Black (UKBBL, *n* = 232), ABCD Hispanic (ABCDH, *n* = 768), and ABCD African American (ABCDA, *n* = 1,257). We meta-analyzed these four non-European GWAS (total *n* = 2,703) and found that 39 (6.4%) passed the Bonferroni significance level and 601 (99.2%) were significant at FDR 5% level. Some associations with rs3781658 (*ANO1*), rs7083220 (*PWWP2B*), rs9373978 (*FHL5*), rs11187838 (*PLCE1*), and rs35124509 (*EPHA3*) were replicated in both European and non-European datasets at the stringent Bonferroni significance level. Moreover, we performed a third meta-analysis to combine all of the eight validation datasets, after which the number of replicated associations moved up to 136 (22.4%) and 602 (99.3%) at Bonferroni and FDR significance levels, respectively. These results are summarized in **Supplementary Table 4**. Overall, our results suggest that the associated genetic loci discovered in UKB British GWAS have high generalizability in independent rsfMRI studies, despite the fact that these studies may use different imaging protocols/MRI scanners and recruit participants from different age groups. The strong homogeneity of GWAS results likely benefit, in part, from the consistent rsfMRI processing procedures that we applied to these datasets.

In addition, we utilized polygenic risk scores^55^ (PRS) derived from UKB British GWAS for further evidence of replication (Methods). For the 197 traits that had significant variants, 168 had significant PRS in at least one of the four European validation GWAS datasets at FDR 5% level (197 × 4 tests, **Supplementary Table 5**), illustrating the significant out-of-sample prediction power of polygenic influences from our discovery GWAS results. The largest incremental R-squared (after adjusting the effects of age, sex, and ten genetic principal components) were observed on the 2nd, 3rd, 4th, and 6th global functional connectivity measures in UKBW and HCP datasets, which were larger than 5% (range = (5.1%, 5.7%), *P* range = (1.1 × 10^−24^, 4 × 10^−13^)). To evaluate the consistency across ancestry, PRS was also constructed on the four non-European validation datasets. UKBA had the best validation performance among the four datasets, with 86 PRS being significant at FDR 5% level (197 × 4 tests, **Supplementary Table 6**). The number of significant PRS was reduced to 59, 39, and 31 in ABCDH, ABCDA, and UKBBL, respectively. In summary, these PRS results illustrate the overall consistency of genetic effects in European cohorts and also show that there may be population specific influences on brain function in other cohorts, though much smaller sample sizes and difficulty in conducting cross ancestry PRS strongly limit the interpretability of these analyses. More efforts are required to identify causal variants associated with functional brain in global diverse populations and perform better cross-population PRS predictions.

### The shared genetic loci with brain-related complex traits and disorders

To evaluate the shared genetic influences between intrinsic brain activity and other complex traits, we carried out association lookups for independent significant variants (and their LD tags, i.e., variants with LD, *r*^2^ ≥ 0.6) detected in UKB British GWAS (Methods). In the NHGRI-EBI GWAS catalog^56^, our results tagged many variants reported for a wide range of complex traits in different trait domains, such as neurological and psychiatric disorders, cognitive performance, education, bone mineral density, sleep, smoking/drinking, brain structure, and anthropometric traits. Below we highlighted colocalizations in a few selected genomic regions.

The index variants rs429358 (*APOE*), rs34404554 (*TOMM40*), rs157582(*TOMM40*), and rs157592 (*APOC1*) in the 19q13.32 region (**Fig. 2a, Supplementary Fig. 4**) had genetic effects on the amplitude of many functional brain regions that were most in the default mode, central executive (i.e., frontoparietal), attention, and visual networks. It is well known that 19q13.32 is a risk locus of Alzheimer’s disease and rs429358 is one of the two variants in the *APOE* ε4 locus. In this region, we tagged variants associated with dementia and decline in mental ability, including Alzheimer’s disease^57–59^, frontotemporal dementia^60^, cerebral amyloid angiopathy^61^, cognitive decline^62^, cognitive impairment test score^63^, as well as many biomarkers of Alzheimer’s disease, such as neurofibrillary tangles^61^, neuritic plaque^61^, cerebral amyloid deposition^64^, cerebrospinal fluid protein levels^63^, and cortical amyloid beta load^65^. Altered amplitude activity has been widely reported in patients of cognitive impairment and Alzheimer’s disease^66,67^. The brain degeneration related to Alzheimer’s disease may begin in the frontoparietal regions^68^ and was associated with dysfunction of multiple RSNs, especially the default mode network^20^. Our findings suggest the shared genetic influences between intrinsic neuronal activity and brain atrophy of Alzheimer’s disease.

**Figure 2:**
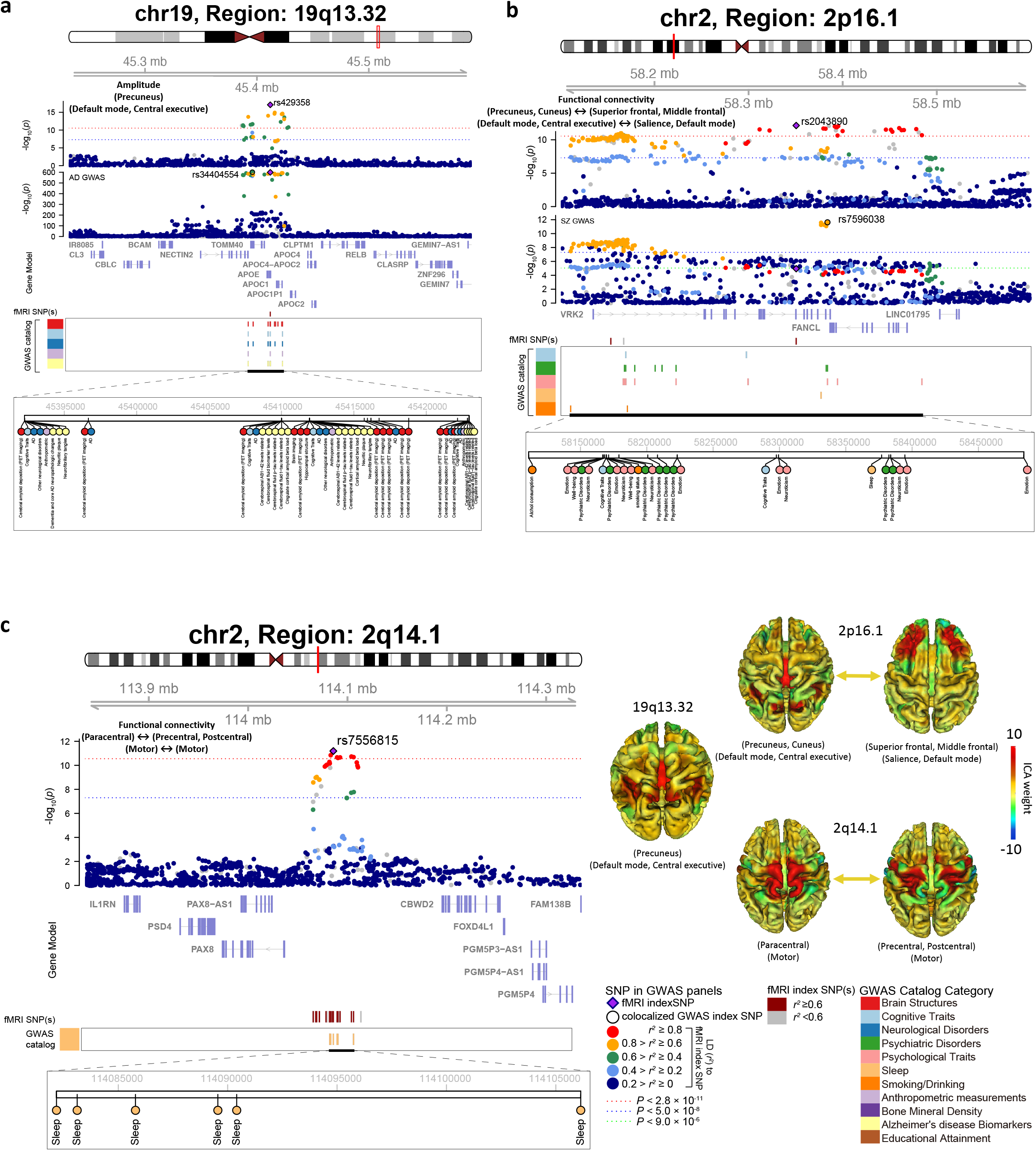
Selected genetic loci that associated with both rsfMRI traits of brain activity and other brain-related complex traits and disorders. We highlight local colocalization (LD *r*^2^ ≥ 0.6) in **a)** 19q13.32 (colocalized with Alzheimer’s disease); **b)** 2p16.1 (with schizophrenia); and **c)** 2q14.1 (with sleep). For example, in 19q13.32, we observed colocalization between the amplitude of the precuneus region in the default mode and central executive networks with Alzheimer’s disease. Location and functional network of the displayed three rsfMRI traits are illustrated on the bottom right. More examples of the shared genetic loci and the involved rsfMRI traits can be found in Supplementary Figures 4-12.

Next, the variant rs62061845 (*KANSL1*) in the 17q21.31 region (**Supplementary Fig. 5**) was associated with functional connectivity over the inferior frontal, middle frontal, superior frontal, middle temporal, and supplementary motor area regions in the default mode and salience networks. Variants in LD with rs62061845 have been frequently reported to be associated with Parkinson’s disease studies^69–73^. As a system-level progressive neurodegenerative disorder^74^, Parkinson’s disease not only leads to motor abnormalities, but also has non-motor symptoms such as temporal perception abnormalities^75^ and impaired connectivity among frontal regions^76^. Cognitive dysfunction and disrupted coupling between default mode and salience networks were commonly reported in Parkinson’s disease^17^. In addition to Parkinson’s disease, the 17q21.31 region was widely related to other complex traits, including neurological disorders (e.g., Alzheimer’s disease^77^, corticobasal degeneration^78^, progressive supranuclear palsy^79^), psychiatric disorders (e.g., autism spectrum disorder^80^, depressive symptoms^81^), educational attainment^82^, psychological traits (e.g., neuroticism^81^), cognitive traits (cognitive ability^83^), sleep^84^, heel bone mineral density^85^, alcohol use disorder^86^, subcortical brain volumes^38^, cortical surface area and thickness^36^, and white matter microstructure^53^.

In addition, the 2p16.1 (**Fig. 2b, Supplementary Fig. 6**) and 5q15 (**Supplementary Fig. 7**) regions were mainly associated with interactions among the central executive, default mode, and salience networks. We observed colocalizations with psychiatric disorders (e.g., schizophrenia^87^, MDD^88^, depressive symptoms^89^, autism spectrum disorder^90^), psychological traits (e.g., neuroticism^81^, well-being spectrum^91^), sleep^92^, cognitive traits (e.g., intelligence^93^), and educational attainment^82^. Dysregulated triple network interactions were frequently reported in patients of schizophrenia^94^, depression^95^, and autism spectrum disorder^96^. Similarly, the 2q24.2 (**Supplementary Fig. 8**) and 10q26.13 (**Supplementary Fig. 9**) regions had genetic effects on functional connectivity traits involved in the central executive, default mode, salience, and limbic networks. In these two regions, our identified variants tagged those that have been implicated with schizophrenia^97^, educational attainment^82^, cognitive traits (e.g., cognitive ability^83^), smoking/drinking (e.g., smoking status^98^, alcohol consumption^99^), hippocampus subfield volumes^100^, and heel bone mineral density^85^. We also observed colocalizations in some other genomic regions, such as in 2q14.1 region (**Fig. 2c, Supplementary Fig. 10**) with sleep traits (e.g., sleep duration^84^, insomnia^92^), in 3p11.1 (**Supplementary Fig. 11**) with cognitive traits (e.g., intelligence^101^, math ability^82^), and in 5q14.3 (**Supplementary Fig. 12**) with cognitive traits^83^ and educational attainment^82^. All of these results are summarized in **Supplementary Table 7**. In summary, intrinsic brain function has wide genetic links to a large number of brain-related complex traits and clinical outcomes, especially neurological and psychiatric disorders and cognitive traits. Integration of GWAS of brain function with these clinical outcomes may help to explain the underlying brain functional mechanisms leading to risk for these disorders.

### Genetic correlations with brain structure, brain disorders, and cognition

The intricate brain neuroanatomical structure is fundamental in supporting brain function. To explore whether genetically mediated brain structural changes were associated with brain function, we examined pairwise genetic correlations (gc) between 1,777 intrinsic brain activity traits and 315 brain structure traits via LDSC^102^ (Methods), including 100 regional brain volumes^37^ and 215 DTI traits of brain structural connectivity in white matter tracts^103^. There were 151 significant pairs between 94 intrinsic brain functional traits and 73 brain structural traits at FDR 5% level (315 × 1,777 tests, |gc| range = (0.22, 0.61), *P* range = (1.2 × 10^−21^, 1.5 × 10^−5^), **Supplementary Table 8**).

We found significant genetic correlations between regional brain volumes and functional connectivity strengths (|gc| range = (0.22, 0.61), *P* range = (1.2 × 10^−21^, 1.2 × 10^−5^), **Supplementary Fig. 13**). Most of the observed correlations were related to higher order brain functional networks, particularly the attention, default mode, salience, and central executive networks. For example, the insula has been widely implicated to be associated with multiple functions, including but not limited to emotion, addiction, and cognition through extensive connections to neocortex, the limbic system, and amygdala^104^. We observed genetic correlations between insula volumes and the connection strengths of multiple pairs of brain regions (|gc| range = (0.22, 0.27), *P* < 1.2 × 10^−5^**, Figs. 3a-b**), which were largely in the default mode and central executive networks, including the angular and the inferior and superior frontal regions. Similarly, left inferior parietal lobule volume exhibited strong genetic correlations with connectivity strengths over multiple pairs of brain regions that were known to be a part of the default mode, visual, attention, and salience networks (|gc| range = (0.34, 0.49), *P* < 9.7 × 10^−6^, **Supplementary Fig. 14a**). Interestingly, however, the above identified genetic correlations appeared to be more specific to the left but not right. The inferior parietal has been implicated to be associated with language function and is connected with the Broca’s region via the superior longitudinal fasciculus (SLF)^105–107^. Considering language processing is left-lateralized in about 95% of right-handers and 75% of left-handers^108–111^, the observed associations of the left inferior parietal are consistent with the results reported in the literature. In addition, we observed spatial colocalizations between regional brain volumes and their genetically correlated functional connectivity traits in multiple brain regions. For instance, left pericalcarine volume was genetically correlated with the connectivity strengths among its neighboring regions, such as the calcarine, superior occipital, cuneus, precuneus, and lingual, which were largely in the visual, default mode, and central executive networks (**Fig. 3c**). More spatial overlap/proximity examples included the associations between right precuneus volume and functional connectivity pairs over the precuneus, angular, inferior parietal, and middle temporal regions (**Supplementary Fig. 14b**); and the associations between postcentral volumes and functional interactions among the postcentral, inferior and superior parietal, supramarginal, and precuneus regions (**Supplementary Fig. 14c**).

**Figure 3:**
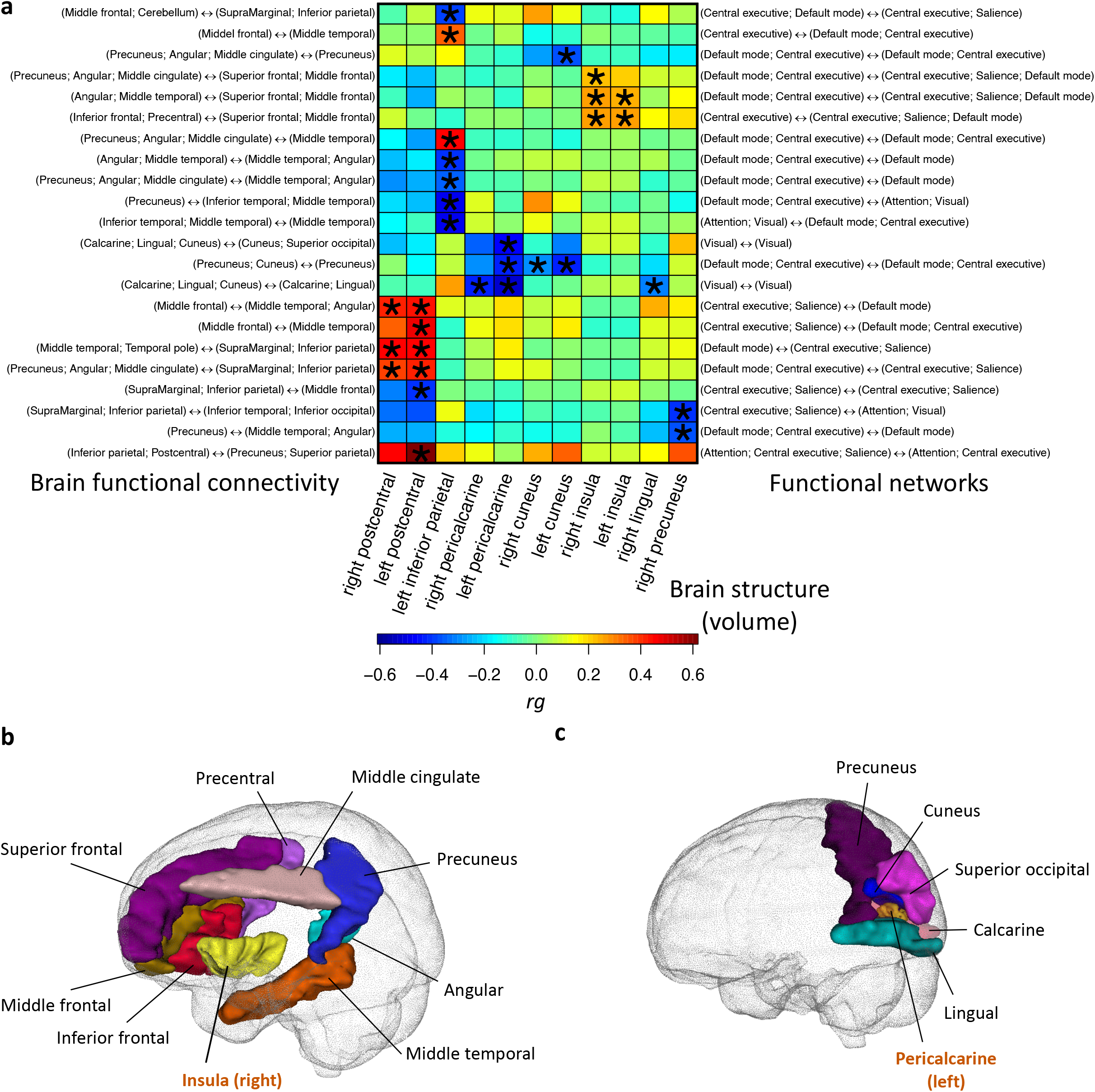
Selected pairwise genetic correlations between functional connectivity traits and regional brain volumes. **a)** The asterisks highlight significant associations after controlling the false discovery rate at 0.05 level. The left y-axis lists the location of functional connectivity traits, the right y-axis shows the associated functional networks, and the x-axis provides the name of regional brain volumes. The colors represent genetic correlations (rg). **b)** Location of the right insula and its neighboring brain regions whose functional connectivity strengths were genetically correlated with the right insula volume. The colors describe different brain regions. **c)** Location of the left pericalcarine and its neighboring brain regions whose functional connectivity strengths were genetically correlated with the left pericalcarine volume.

Significant genetic correlations were also observed between brain structural connectivity and functional connectivity (|gc| range = (0.25, 0.49), *P* range = (5.5 × 10^−10^, 1.3 × 10^−5^), **Supplementary Fig. 15**). Many of the white matter tracts, in particular the SLF and corpus callosum, manifested a strong genetic correlation with the interactions of functional networks (**Fig. 4a**). These results provided genetic evidence on how these distributed networks communicate across large distances. The SLF has been widely documented connecting brain regions in temporal, parietal, and frontal lobes^112^. Functionally, SLF has been reported associated with a wide array of brain functions, including working memory^113^, attention^114,115^, and language functions^116,117^. We observed significant genetic correlations between SLF and connectivity strengths over multiple pairs of brain regions including the frontal, parietal, and temporal regions (|gc| range = (0.33, 0.49), *P* < 2.4 × 10^−6^, **Fig. 4b**). For example, a significant association between insula and temporal connection and SLF was observed. This finding is consistent with the well documented broad functions of insula, including attention and salience processes^104^. Furthermore, parietal and frontal connections most likely reflected attention and executive control networks. Moreover, the splenium of corpus callosum (SCC) is located in the most posterior part of the corpus callosum and connects brain regions in the temporal, posterior parietal, and occipital lobes. Our results show that SCC was genetically associated with brain regions within the parietal lobe (|gc| range = (0.34, 0.48), *P* < 6.5 × 10^−6^, **Fig. 4c**). In particular, multiple regions connected to the precuneus were observed, such as the inferior parietal, supramarginal, and occipital regions. The precuneus has been shown to connect multiple cortical and subcortical regions. Functionally, the precuneus is one of the critical areas of the default mode network and has also been implicated to be associated with attention as well as memory functions^118^. Our findings suggest that these connections may be genetically mediated by the SCC. Besides functional connectivity traits, amplitude traits also had significant genetic associations with regional brain volumes and white matter tracts (**Supplementary Figs. 16-17**, **Supplementary Note**). Overall, our results uncover the genetic links between intrinsic brain function networks and the associated structural substrates. As illustrated, a few pairs of the genetically correlated brain functional and structural traits show high congruity in spatial location and the involved functions. There has been growing interest to understand how brain topography interacts with brain functional networks^119^. To our knowledge, our results are the first to indicate that genetic changes in brain structure may also impact brain function.

**Figure 4:**
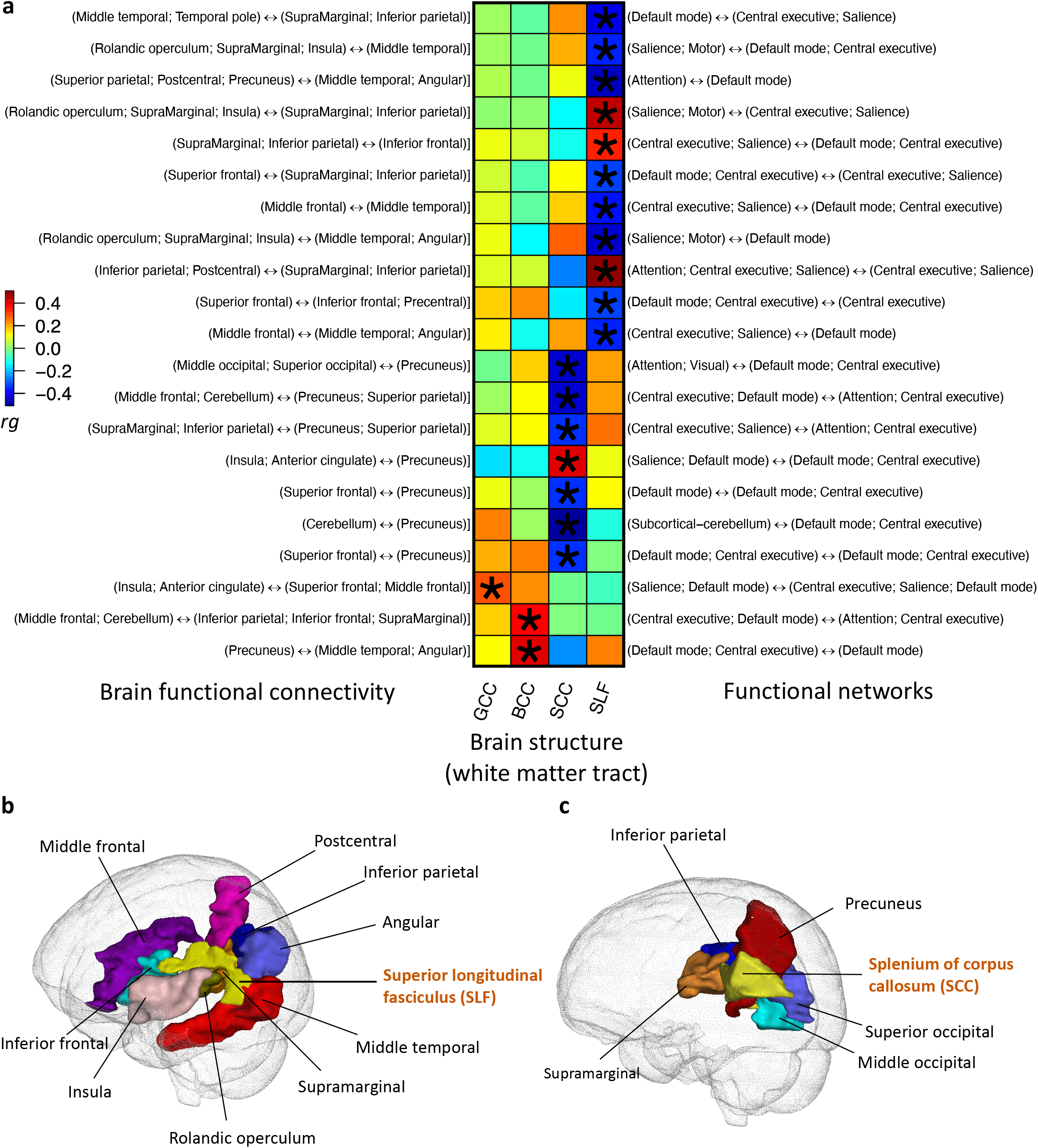
Selected pairwise genetic correlations between functional connectivity traits and fractional anisotropy (FA) of white matter tracts. **a)** The asterisks highlight significant associations after controlling the false discovery rate at 0.05 level. The left y-axis lists the location of functional connectivity traits, the right y-axis shows the associated functional networks, and the x-axis provides the name of white matter tracts. The colors represent genetic correlations (rg). **b)** Location of the SLF (left part) and its neighboring brain regions whose functional connectivity strengths were genetically correlated with the FA of SLF. The colors describe different brain regions. **c)** Location of the SCC (left part) and its neighboring brain regions whose functional connectivity strengths were genetically correlated with the FA of SCC.

Next, we examined the genetic correlations between 1,777 intrinsic brain activity traits and 30 other complex traits, mainly focusing on brain disorders and cognition (**Supplementary Table 9**). We found 176 significant pairs between 26 complex traits and 102 intrinsic brain activity traits at FDR 5% level (30 × 1,777 tests, *P* range = (8.6 × 10^−12^, 2.3 × 10^−3^), **Supplementary Table 10**). Particularly, functional connectivity strengths were genetically correlated with a few brain disorders, including attention deficit hyperactivity disorder (ADHD), schizophrenia (SCZ), major depressive disorder (MDD), and cross disorder (five major psychiatric disorders^120^) (|gc| range = (0.18, 0.37), *P* < 1.2 × 10^−4^, **Fig. 5a**). For example, we observed a significant genetic correlation between ADHD and functional interactions among the precentral, supplementary motor area, superior frontal, putamen, and caudate regions, which were largely in the attention, salience, motor, and subcortical-cerebellum networks (**Fig. 5b**). These brain regions have been widely implicated with ADHD in previous studies. ADHD patients have been observed to have stronger connectivity across the supplementary motor area, precentral, and superior frontal regions^121^. These regions are also associated with difficulties in performing some fine motor skills^122^. In addition, the putamen and caudate regions compose the dorsal striatum, one largest part of the basal ganglia, which is important in controlling motor functions^123,124^. Moreover, significant genetic correlations were observed between SCZ and connection strengths over the precentral, postcentral, precuneus, frontal, and superior parietal regions (**Fig. 5c**); and between MDD and the interactions among the middle temporal, angular, and superior and middle frontal regions (**Fig. 5d**, **Supplementary Note**).

**Figure 5:**
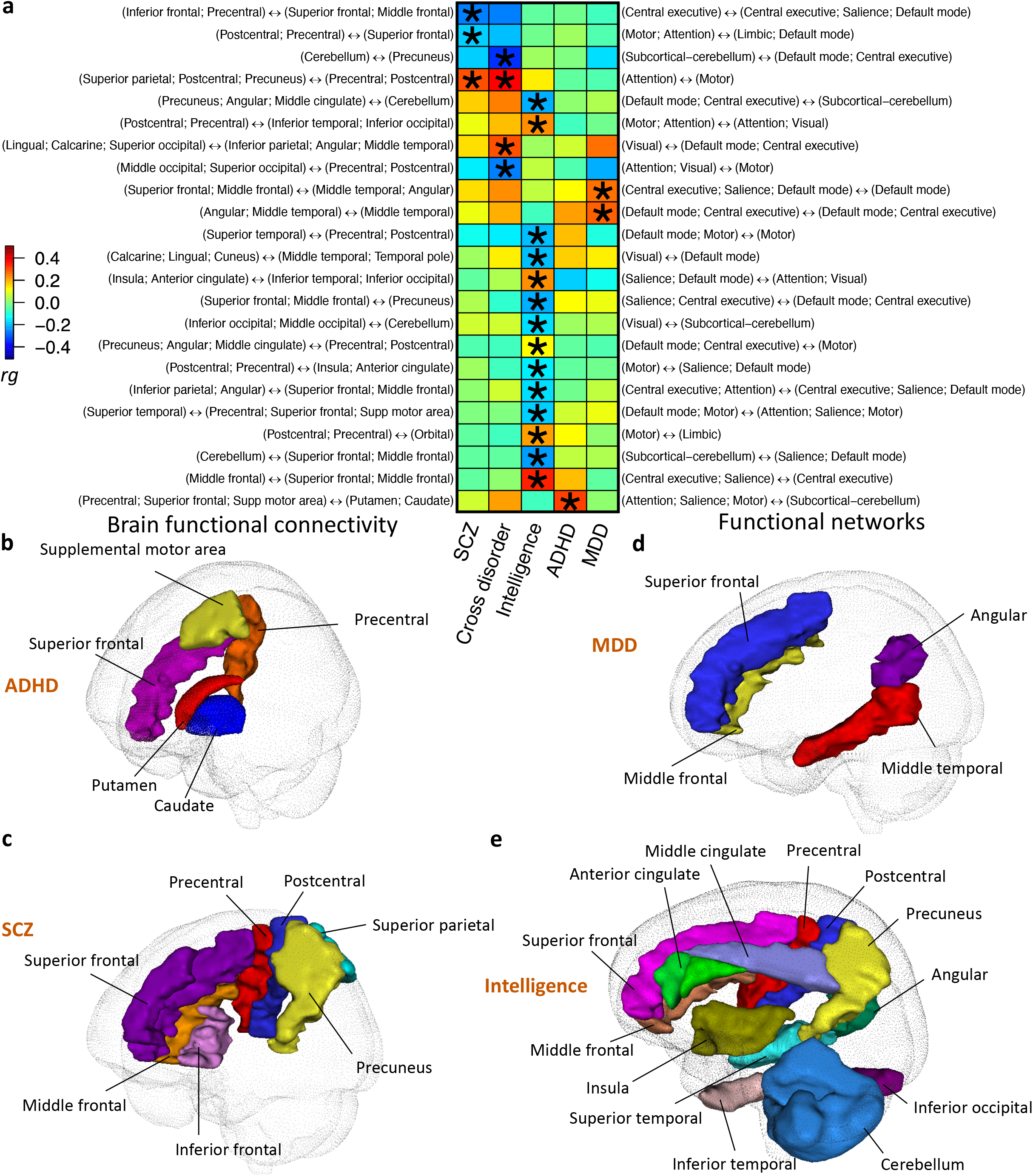
Selected pairwise genetic correlations between functional connectivity traits and other brain-related traits/disorders. **a)** The asterisks highlight significant associations after controlling the false discovery rate at 0.05 level. The left y-axis lists the location of functional connectivity traits, the right y-axis shows the associated functional networks, and the x-axis provides the name of other brain-related traits/disorders. The colors represent genetic correlations (rg). **b-e)** Location of the brain regions whose functional connectivity strengths were genetically correlated with **b)** attention-deficit/hyperactivity disorder (ADHD); **c)** schizophrenia (SCZ); **d)** major depressive disorder (MDD); and **e)** intelligence. The colors describe different brain regions.

In addition, many genetic correlations were observed between functional connectivity and cognitive traits studied in previous GWAS, including intelligence, cognitive performance, general cognitive function, and numerical reasoning. For example, intelligence had genetic correlations with connection strengths over multiple brain regions (|gc| range = (0.11, 0.34), *P* < 1.8 × 10^−4^, **Fig. 5e**). The strongest correlation located at the superior and middle frontal regions in the central executive and salience networks. It is known that the frontal lobe is associated with higher level cognitive skills, such as problem solving, thinking, planning, and organizing^125^. Wang, et al. ^126^ revealed a general intelligence network for logical-math, general intelligence, and linguistic skills, which widely included frontal, parietal, occipital, temporal, and limbic regions. Furthermore, significant genetic correlations were broadly observed on subjective well-being, education, neuroticism, sleep, risk tolerance, automobile speeding, manual occupation, BMI, high blood pressure, and behavioral factors (drinking and smoking) (**Supplementary Figs. 18-19**). More details and interpretations can be found in **Supplementary Note**.

### Gene-level association analysis and biological annotations

Gene-level association was tested via MAGMA^127^ (Methods), which detected 970 significant gene-trait associations (*P* < 1.5 × 10^−9^, adjusted for 1,777 phenotypes) for 123 genes (**Supplementary Fig. 20, Supplementary Table 11**). In addition, we applied FUMA^54^ to map significant variants (*P* < 2.8 × 10^−11^) to genes via physical position, expression quantitative trait loci (eQTL) association, and 3D chromatin (Hi-C) interaction, which yielded 197 more associated genes that were not discovered in MAGMA (276 in total, **Supplementary Table 12**). For the 320 genes associated with intrinsic brain activity in either MAGMA or FUMA, 84 had been linked to white matter microstructure^103^, 48 were reported to be associated with regional brain volumes^37^, and 42 were related to both of them (**Supplementary Table 13**). These triple overlapped genes were also widely associated with other complex traits, such as Parkinson’s disease, neuroticism, stroke, alopecia, handedness, and intelligence (**Supplementary Table 14**), providing more insights into the genetic overlaps among brain structure, brain function, and other brain-related traits. For example, *MAPT, NSF, WNT3*, and *LRRC37A3* were risk genes of Parkinson’s disease, which were also associated with pallidum volumes^37^, white matter microstructure^103^, and intrinsic functional connectivity in central executive, default mode, and salience networks. These complementary neuroimaging traits had all been used to study the pathophysiology of Parkinson’s disease^128–130^. Similarly, *CDKN2C* and *FAF1* were associated with ischemic stroke^131^ as well as multiple neuroimaging traits of brain structure and function. In addition, 4 of our intrinsic brain activity associated genes (*CALY, SLC47A1, CYP2C8*, and *CYP2C9*) were targets for 11 nervous system drugs^132^, such as 4 psycholeptics (ATC code: N05) to produce calming effects, 2 anti-depressants (N06A) to treat MDD and related conditions, 2 anti-migraine (N02C), and one anti-dementia (N06D) (**Supplementary Table 15**).

It is of particular interest to study the functional connectivity dysfunction in Alzheimer’s disease and identify the overlapped genes^20,133^. Our gene-level analysis replicated *APOE* and *SORL1*, which were frequently targeted in Alzheimer’s disease-candidate gene studies of functional connectivity^23,134^. More importantly, we uncovered more overlapped genes between intrinsic brain activity and Alzheimer’s disease, such as *PVRL2, TOMM40, APOC1, MAPK7, CLPTM1, HESX1, BCAR3, ANO3*, and *YAP1* (**Supplementary Table 16**). Interestingly, through the BIG-KP server, we found that these genes had much stronger associations with intrinsic brain function than brain structure. We also observed many pleiotropic genes associated with serum metabolite, low density lipoprotein cholesterol, high density lipoprotein cholesterol, triglyceride, type II diabetes mellitus, and blood protein measurements, all of which might be related to the Alzheimer’s disease^135,136^. These results expand the overview of the shared genetic components among metabolic dysfunction, blood biomarkers, brain function in Alzheimer’s disease research, suggesting the potential value of integrating these traits in future studies.

To identify the tissues and cell types in which genetic variation yields differences in brain functional connectivity, we performed partitioned heritability analyses^137^ for tissue type and cell type specific regulatory elements^138^ (Methods). We focused on the 10 functional connectivity traits that had heritability higher than 30%. At FDR 5% level, the most significant enrichments of heritability were observed in active gene regulation regions of fetal brain tissues, neurospheres, and neuron/neuronal progenitor cultured cells (**Supplementary Fig. 21, Supplementary Table 17**). We also tried to further identify brain cell type specific enrichments using chromatin accessibility data of two main gross brain cell types^139^ (i.e., neurons (NeuN+) and glia (NeuN-)) and multiple neuronal and glial cell subtypes, including oligodendrocyte (NeuN-/Sox10+), microglia, and astrocyte (NeuN-/Sox10-), as well as GABAergic (NeuN+/Sox6+) and glutamatergic neurons (NeuN+/Sox6-). Although enrichments were observed in some cell types, few of them remained significant after adjusting for multiple testing (**Supplementary Fig. 22, Supplementary Table 18**). Next, we performed MAGMA tissue-specific gene property^127^ analysis for 13 GTEx^140^ (v8) brain tissues (Methods). We found that genes with higher expression levels in human brain tissues generally had stronger associations with intrinsic brain activity, particularly for tissues sampled from cerebellar hemisphere and cerebellum regions (*P* < 1.9 × 10^−5^, **Supplementary Fig. 23**, **Supplementary Table 19**).

Among the associated variants of intrinsic brain activity, a few resided in frequently interacting regions (FIREs) and topologically associating domain (TAD) boundaries in brain tissues^141,142^ (**Supplementary Table 20**). Partitioned heritability analysis also provided suggestive evidence of heritability enrichment in these FIREs and TAD boundaries (**Supplementary Fig. 24**, **Supplementary Table 21**). We performed additional gene mapping using 14 recent Hi-C datasets of brain tissue and cell types^141–145^ (Methods). This Hi-C gene mapping prioritized 29 genes, 14 of which were not identified by the Hi-C analysis in FUMA^54^ (**Supplementary Table 22**). Many of the newly mapped genes have been reported for brain-related disorders/conditions, sleep, and intelligence, including *APOE, HSPG2, APOC1, UFL1, NR2F1, NPM1, FAM172A, FADD, FHL5*, and *EPHA3*. Finally, MAGMA^127^ gene-set analysis was performed to prioritize the enriched biological pathways (Methods). We found 59 significantly enriched gene sets after Bonferroni adjustment (*P* < 1.8 × 10^−9^, **Supplementary Table 23**). Multiple pathways related to nervous system were detected, such as “go neurogenesis” (GO: 0022008), “go neuron differentiation” (GO: 0030182), “go regulation of nervous system development” (GO: 0051960), “go regulation of neuron differentiation” (GO: 0045664), “go cell morphogenesis involved in neuron differentiation” (GO: 0048667), and “go neuron development” (GO: 0048666).

## DISCUSSION

In the present study, we evaluated the influences of common variants on intrinsic brain functional architecture using harmonized rsfMRI data of 44,190 subjects from four independent studies. Genome-wide association analysis found hundreds of novel loci related to intrinsic brain activity in the UKB British cohort, which were successfully replicated in independent datasets. The interactions across core neurocognitive networks (central executive, default mode, and salience) in the triple network model had genetic links with cognition and multiple brain disorders. Shared genetic influences among functional, structural, and diffusion neuroimaging traits were also uncovered, showing that brain structure and function are intimately related. Gene-level analysis detected many overlapped genes between intrinsic brain activity and Alzheimer’s disease. We also detected a colocalization between one of the two variants in the *APOE* ε4 locus and function of the default mode, central executive, attention, and visual networks, which may explain in part the functional mechanism underlying Alzheimer’s risk. The enriched tissues and biological pathways were also prioritized in bioinformatic analyses. Compared to the previous study^31^ with about 8,000 subjects, this large-scale GWAS much improved our understanding of the genetic architecture of functional human brain.

Our study faces a few limitations. First, the samples in our discovery GWAS were mainly from European ancestry. In our PRS analysis, we illustrated a relatively poor replication of the European GWAS results within validation cohorts with non-European ancestry. The non-European GWAS was of small sample size, so population specific influences will be better understood when more data from global populations become available. Second, our study focused on the brain functional activity at rest. A recent study^28^ had found that combining rsfMRI and task functional magnetic resonance imaging (tfMRI) may result in higher heritability estimates and potentially boost the GWAS power. Thus, future studies could model rsfMRI and tfMRI together to uncover more insights into the genetic influences on brain function. In addition, we applied ICA in this study, which was a popular approach to characterize the functionally connected brain^6^. It is also of great interest to evaluate the performance of other popular rsfMRI approaches (such as seed-based analysis) in these large-scale datasets. Finally, although we found genetic links between brain function and other complex traits, future work is needed to dissect the underlying mechanisms by which genetic variation leads to differences in brain activity. We expect that accumulating publicly available imaging genetics data resources will lead to a better understanding of specific genes involved in human brain structure function relationships and how variants can alter these relationships leading to risk for neuropsychiatric disorders.

## URLs

Brain Imaging Genetics Knowledge Portal (BIG-KP), https://bigkp.org/;

Brain Imaging GWAS Summary Statistics, https://github.com/BIG-S2/GWAS;

UKB Imaging Pipeline, https://git.fmrib.ox.ac.uk/falmagro/UK_biobank_pipeline_v_1;

PLINK, https://www.cog-genomics.org/plink2/;

GCTA & fastGWA, http://cnsgenomics.com/software/gcta/;

METAL, https://genome.sph.umich.edu/wiki/METAL;

FUMA, http://fuma.ctglab.nl/;

MGAMA, https://ctg.cncr.nl/software/magma;

LDSC, https://github.com/bulik/ldsc/;

FINDOR, https://github.com/gkichaev/FINDOR;

NHGRI-EBI GWAS Catalog, https://www.ebi.ac.uk/gwas/home;

The atlas of GWAS Summary Statistics, http://atlas.ctglab.nl/.

## METHODS

Methods are available in the ***Methods*** section.

*Note: One supplementary information pdf file and one supplementary table zip file are available*.

## ACKNOWLEDGEMENTS

This research was partially supported by U.S. NIH grants MH086633 (H.Z.) and MH116527 (TF.L.). We thank the individuals represented in the UK Biobank, ABCD, HCP, and PNC studies for their participation and the research teams for their work in collecting, processing and disseminating these datasets for analysis. We gratefully acknowledge all the studies and databases that made GWAS summary data available. This research has been conducted using the UK Biobank resource (application number 22783), subject to a data transfer agreement. Part of the data used in the preparation of this article were obtained from the Adolescent Brain Cognitive Development (ABCD) Study (https://abcdstudy.org), held in the NIMH Data Archive (NDA). This is a multisite, longitudinal study designed to recruit more than 10,000 children age 9-10 and follow them over 10 years into early adulthood. The ABCD Study is supported by the National Institutes of Health and additional federal partners under award numbers U01DA041022, U01DA041028, U01DA041048, U01DA041089, U01DA041106, U01DA041117, U01DA041120, U01DA041134, U01DA041148, U01DA041156, U01DA041174, U24DA041123, U24DA041147, U01DA041093, and U01DA041025. A full list of supporters is available at https://abcdstudy.org/federal-partners.html. A listing of participating sites and a complete listing of the study investigators can be found at https://abcdstudy.org/scientists/workgroups/. ABCD consortium investigators designed and implemented the study and/or provided data but did not necessarily participate in analysis or writing of this report. This manuscript reflects the views of the authors and may not reflect the opinions or views of the NIH or ABCD consortium investigators. Support for the collection of the PNC datasets was provided by grant RC2MH089983 awarded to Raquel Gur and RC2MH089924 awarded to Hakon Hakonarson. All PNC subjects were recruited through the Center for Applied Genomics at The Children’s Hospital in Philadelphia. HCP data were provided by the Human Connectome Project, WU-Minn Consortium (Principal Investigators: David Van Essen and Kamil Ugurbil; 1U54MH091657) funded by the 16 NIH Institutes and Centers that support the NIH Blueprint for Neuroscience Research; and by the McDonnell Center for Systems Neuroscience at Washington University.

## AUTHOR CONTRIBUTIONS

B.Z., H.Z., J.L.S., S.M.S., and Y.L. designed the study. B.Z., TF.L., D.X., X.W., Y.Y., TY.L., N.M., Q.S., YC.Y. analyzed the data. TF. L., Z.Z., and Y.S. downloaded the datasets, processed rsfMRI data, and undertook quantity controls. P.R., M.E.H., J.B., and J.F.F. analyzed brain cell chromatin accessibility data. B.Z. and H.Z. wrote the manuscript with feedback from all authors.

**CORRESPINDENCE AND REQUESTS FOR MATERIALS** should be addressed to H.Z.

## COMPETETING FINANCIAL INTERESTS

The authors declare no competing financial interests.

## METHODS

### Imaging phenotypes and datasets

The rsfMRI datasets were consistently processed following the procedures in UK Biobank imaging pipeline^9^. Details about image acquisition, preprocessing, and phenotype generation in each dataset can be found in **Supplementary Note**. Following the previous study^31^, we generated two groups of phenotypes, including 76 node amplitude traits reflecting the spontaneous neuronal activity, and 1,695 pairwise functional connectivity traits quantifying co-activity for node pairs, as well as 6 global functional connectivity measures to summarize all pairwise functional connectivity. To aid interpretation of these phenotypes, the functional brain regions characterized in ICA were labelled using the automated anatomical labeling atlas^51^ (**Supplementary Table 24**) and were mapped onto major functional networks defined in Yeo, et al. ^14^ and Finn, et al. ^12^ (**Supplementary Figs. 25-26**). The assigned location and functional networks are provided in **Supplementary Table 25**. Details of our mapping procedures are provided in **Supplementary Note**. For each continuous phenotype or covariate variable, values greater than five times the median absolute deviation from the median value were removed. We analyzed the following nine datasets separately: 1) the UKB discovery GWAS, which used data of individuals of British ancestry^146^ in the UKB study (*n* = 34,691); 2) four European validation GWAS: UKB White but Non-British (UKBW, *n* = 1,970), ABCD European (ABCDE, *n* = 3,821), HCP (*n* = 495), and PNC (*n* = 510); 3) two non-European UKB validation GWAS: UKB Asian (UKBA, *n* = 446) and UKB Black (UKBBL, *n* = 232); and 4) two non-European non-UKB validation GWAS, including ABCD Hispanic (ABCDH, *n* = 768) and ABCD African American (ABCDA, *n* = 1,257). See **Supplementary Table 26** for a summary of these datasets and demographic information. The assignment of ancestry in UKB was based on self-reported ethnicity (Data-Field 21000), which was verified in Bycroft, et al. ^146^. The ancestry in ABCD was assigned by combining the self-reported ethnicity and ancestry inference results as in Zhao, et al. ^103^.

### GWAS discovery and validation

Details of genotyping and quality controls can be found in **Supplementary Note**. SNP heritability was estimated by GCTA^52^ using all autosomal SNPs in the UKB British cohort. We adjusted the effects of age (at imaging), age-squared, sex, age-sex interaction, age-squared-sex interaction, imaging site, and the top 40 genetic principle components (PCs). Genome-wide association analysis was performed in linear mixed effect model using fastGWA^147^, while adjusting the same set of covariates as in GCTA. GWAS were also separately performed via Plink^148^ in the eight validation datasets, including UKBW, UKBBL, UKBA, ABCDA, ABCDH, ABCDE, HCP, and PNC, where the effects of age, age-squared, sex, imaging sites (if applicable), scanners (if applicable), age-sex interaction, age-squared-sex interaction, and top ten genetic PCs were adjusted.

To validate results in the UKB British discovery GWAS, meta-analysis was performed using the sample-size weighted approach via METAL^149^. We examined whether the locus-level associations detected in the British GWAS can be replicated in the 1) meta-analyzed four European validation GWAS (UKBW, ABCDE, HCP, and PNC); 2) meta-analyzed four non-European validation GWAS (UKBBL, UKBA, ABCDA, and ABCDH); and 3) the combination of the above eight validation GWAS. Specifically, for each meta-analyzed GWAS, we checked and reported the smallest *P*-value among the variants within each associated locus identified in the UKB British discovery GWAS. Polygenic risk scores (PRS) were constructed on eight validation datasets using Plink. The BLUP effect sizes estimated from GCTA-GREML analysis in UKB British discovery GWAS were used as weights in PRS construction, which accounted for the LD structures. Ambiguous variants (i.e. variants with complementary alleles) were removed from analysis. We tried 17 *P*-value thresholds for variant selection according to their marginal *P*-values from fastGWA: 1, 0.8, 0.5, 0.4, 0.3, 0.2, 0.1, 0.08, 0.05, 0.02, 0.01, 1 × 10^−3^, 1 × 10^−4^, 1 × 10^−5^, 1 × 10^−6^, 1× 10^−7^, and 1 × 10^−8^. The best prediction accuracy achieved by a single threshold was reported for each phenotype, which was measured by the additional phenotypic variation that can be explained by the polygenic profile (i.e., the incremental R-squared), while adjusting for the effects of age, sex, and top ten genetic PCs.

### The shared loci and genetic correlation

The genomic loci associated with intrinsic brain activity traits were defined using FUMA (version 1.3.5e). We input UKB British discovery summary statistics after reweighting the *P*-values using functional information via FINDOR^98^. To define the LD boundaries, FUMA identified independent significant variants, which were defined as variants with a *P*-value smaller than the predefined threshold and were independent of other significant variants (LD *r*^2^ < 0.6). FUMA then constructed LD blocks for these independent significant variants by tagging all variants in LD (*r*^2^ ≥ 0.6) with at least one independent significant variant and had a MAF ≥ 0.0005. These variants included those from the 1000 Genomes reference panel that may not have been included in the GWAS. Moreover, within these significant variants, independent lead variants were identified as those that were independent from each other (LD *r*^2^ < 0.1). If LD blocks of independent significant variants were close (<250 kb based on the closest boundary variants of LD blocks), they were merged into a single genomic locus. Thus, each genomic locus could contain multiple significant variants and lead variants. Independent significant variants and all the variants in LD with them (*r*^2^ ≥ 0.6) were searched by FUMA on the NHGRI-EBI GWAS catalog (version 2019-09-24) to look for previously reported associations (*P* < 9 × 10^−6^) with any traits. LDSC^102^ software (version 1.0.1) was used to estimate and test the pairwise genetic correlation. We used the pre-calculated LD scores provided by LDSC, which were computed using 1000 Genomes European data. We used HapMap3^150^ variants and removed all variants in the major histocompatibility complex (MHC) region. The summary statistics of intrinsic brain activity traits were from the UKB British discovery GWAS and the resources of other summary statistics were provided in **Supplementary Table 9**.

### Gene-level analysis and biological annotation

Gene-based association analysis was performed in UKB British participants for 18,796 protein-coding genes using MAGMA^127^ (version 1.07). Default MAGMA settings were used with zero window size around each gene. We then carried out FUMA functional annotation and mapping analysis, in which variants were annotated with their biological functionality and then were linked to 35,808 candidate genes by a combination of positional, eQTL, and 3D chromatin interaction mappings. Brain-related tissues/cells were selected in all options and default values were used for all other parameters in FUMA. For the detected genes in MAGMA and FUMA, we performed lookups in the NHGRI-EBI GWAS catalog (version 2020-02-08) to explore their previously reported gene-trait associations. We performed heritability enrichment analysis via partitioned LDSC^137^. Baseline models were adjusted when estimating and testing the enrichment scores for our tissue type and cell type specific annotations. Methods to analysis chromatin data of glial and neuronal cell subtypes can be found in Zhao, et al. ^103^. We also performed gene property analysis for the 13 GTEx^140^ v8 brain tissues via MAGMA. Specifically, we examined whether the tissue-specific gene expression levels can be linked to the strength of the gene-trait association. MAGMA was also used to explore the enriched biological pathways, in which we tested 500 curated gene sets and 9,996 Gene Ontology (GO) terms from the Molecular Signatures Database^151^ (MSigDB, version 7.0). Additional gene mapping was performed using 14 Hi-C datasets of brain tissue and cell types from five recent studies, including 1) the promoter capture Hi-C (PCHi-C) data of hippocampus and dorsolateral prefrontal cortex (DLPFC)^143^; 2) the Hi-C data of hippocampus and DLPFC^141^; 3) the Hi-C data from fetal and adult cortices^142^, restricting to the high confidence interactions; 4) the PCHi-C data of primary astrocytes and three types of induced pluripotent stem cell (iPSC)-derived neurons^144^ (cortical, hippocampal, and motor); and 5) proximity ligation assisted chromatin immunoprecipitation (PLAC-seq) data on sorted fetal neuron cells^145^, including radial glial cells, intermediate progenitor cells, neurons, and interneurons. For interaction intensity cutoffs, we used 2 for the −log10(*P*) used in datasets of Jung, et al. ^143^, 0.05 for the *q*-value in Schmitt, et al. ^141^ and Giusti-Rodriguez and Sullivan ^142^, 5 for the Chicago score in Song, et al. ^144^, and 0.01 for the FDR in Song, et al. ^145^.

## Code availability

We made use of publicly available software and tools listed in URLs. Other codes used in our analyses are available upon reasonable request.

## Data availability

Our GWAS summary statistics can be downloaded at https://github.com/BIG-S2/GWAS. The individual-level data used in the present study can be obtained from four publicly accessible data resources: UK Biobank (http://www.ukbiobank.ac.uk/resources/), ABCD (https://abcdstudy.org/), HCP (https://www.humanconnectome.org/), and PNC (https://www.med.upenn.edu/bbl/philadelphianeurodevelopmentalcohort.html). Our results can also be easily browsed through our knowledge portal https://bigkp.org/.

