## Supplementary material for "Common variants contribute to intrinsic human brain functional networks": supp_figures

September 10, 2020

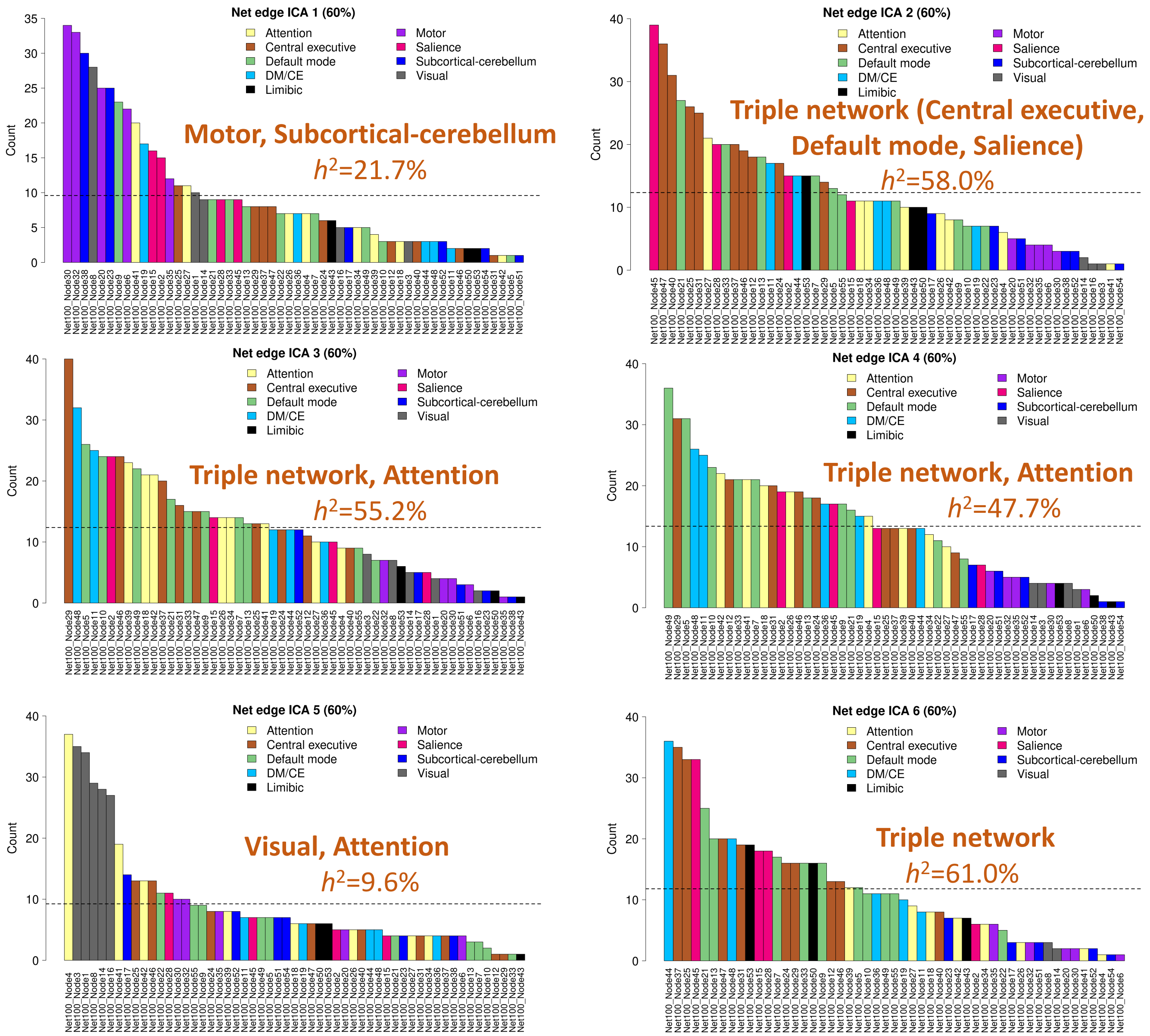


**Supplementary Figure 1:** The top ranked nodes (i.e., ICA components) in each of the 6 global functional connectivity measures and their most related functional networks. The nodes were ranked by their frequencies (counts) among the set of pairwise functional connectivity traits that explain 60% variation of the global functional connectivity measure. The dashed line represents the averaged frequency. The x axis displays the node IDs, the y axis shows their frequencies (counts), and we also label the SNP heritability ($h^2$) of each global functional connectivity measure. Attention, attention network; Motor, motor network; Central executive, central executive network; Salience, salience network; Default mode, default mode network; Subcortical-cerebellum, subcortical-cerebellum network, DM/CE, both default mode and central executive networks; Limbic, limbic network; Visual, visual network. Triple network includes the default mode, central executive, and salience networks.

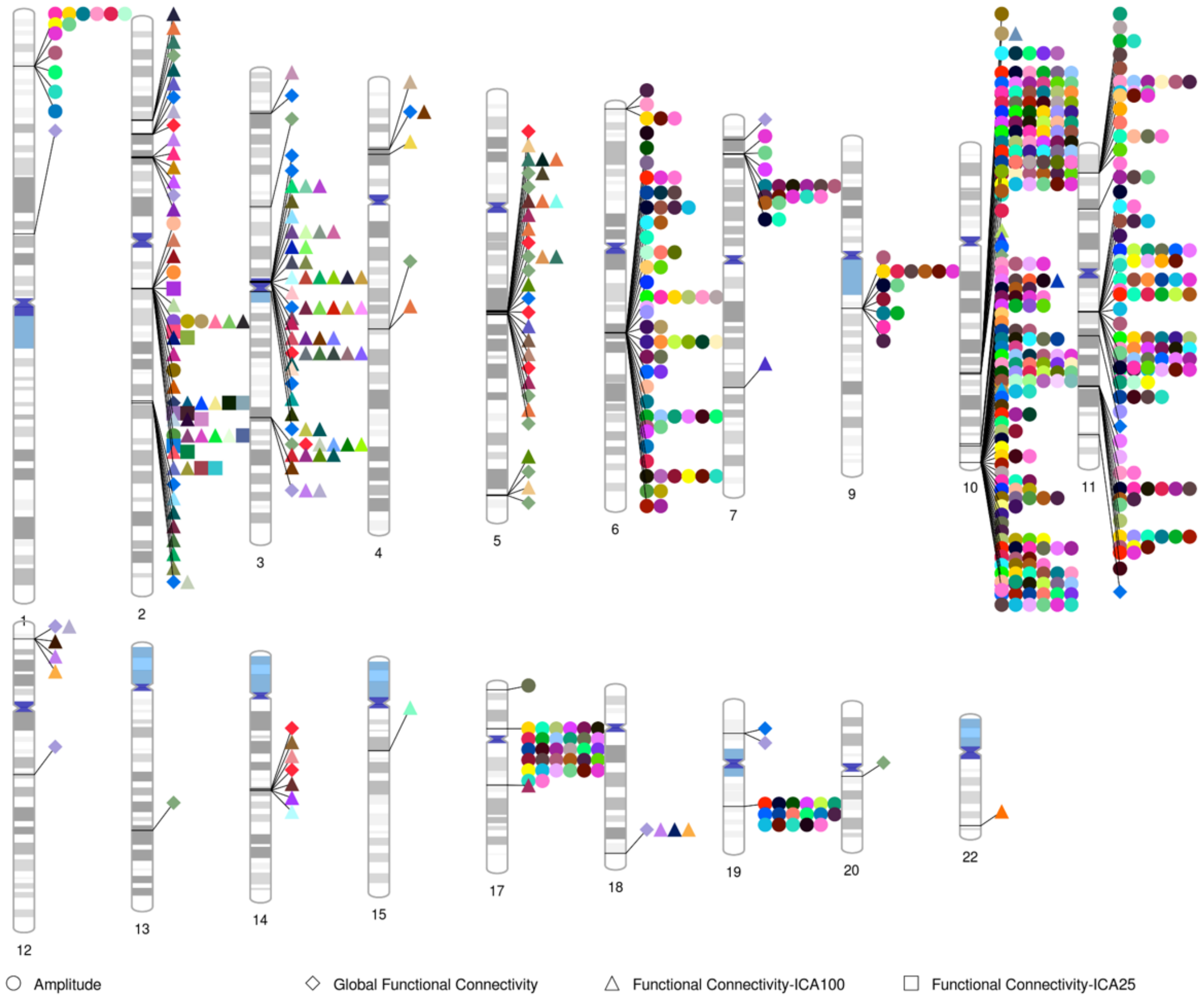


**Supplementary Figure 2:** Ideogram of the loci influencing rsfMRI traits of intrinsic brain activity at the significance level $2.8 \times 10^{-11}$ (i.e., $5 \times 10^{-8}/1,777$). The color represents different rsfMRI traits. The shape of label represents the categories of traits (amplitude, global functional connectivity, pairwise functional connectivity with parcellation of 100 dimensionality, and pairwise functional connectivity with parcellation of 25 dimensionality).

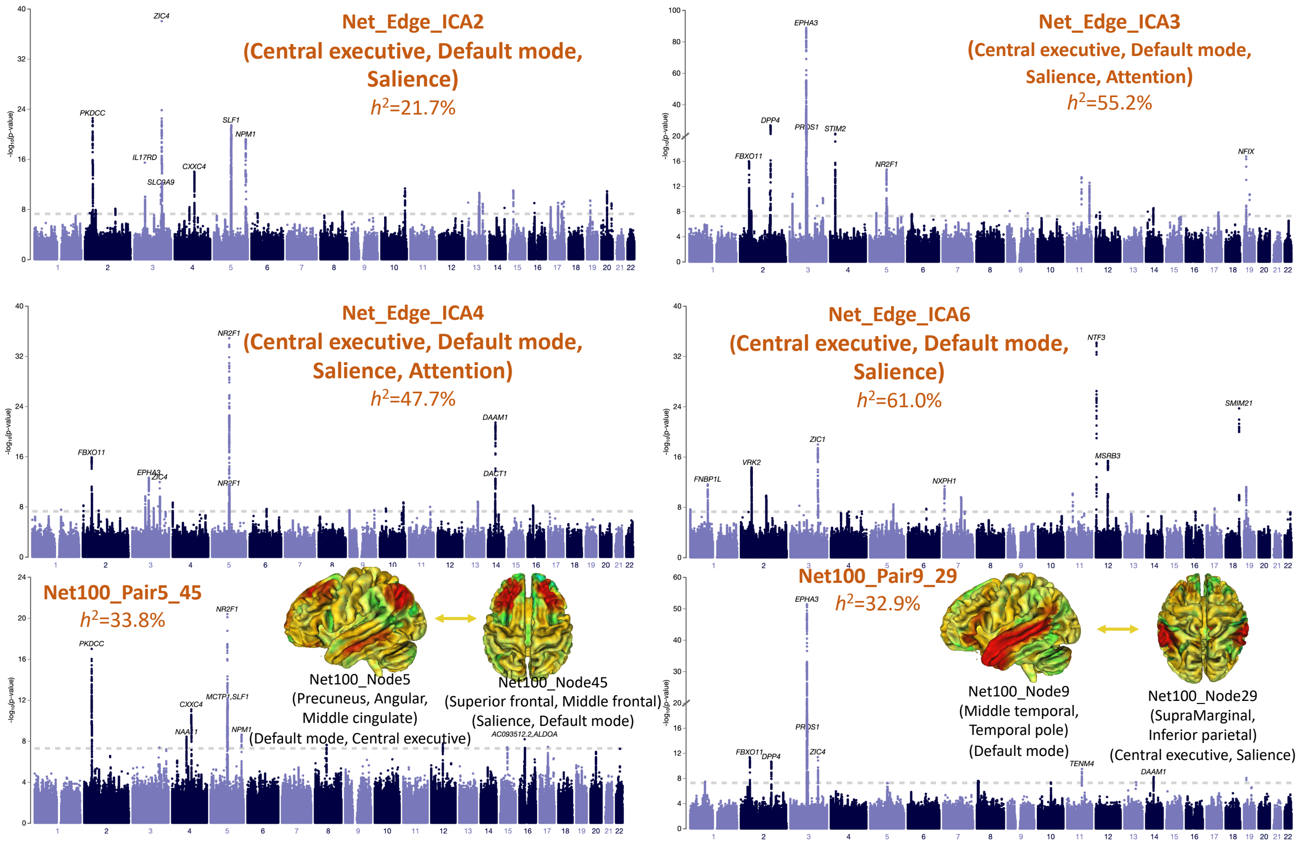


**Supplementary Figure 3:** Manhattan plot of the 6 (4 global and 2 pairwise) functional connectivity traits that had at least 5 independent significant lead variants. For the 4 global functional connectivity measures, we label their most related functional networks as in Supplementary Figure 1. For the 2 pairwise functional connectivity traits, we label their location and functional networks.

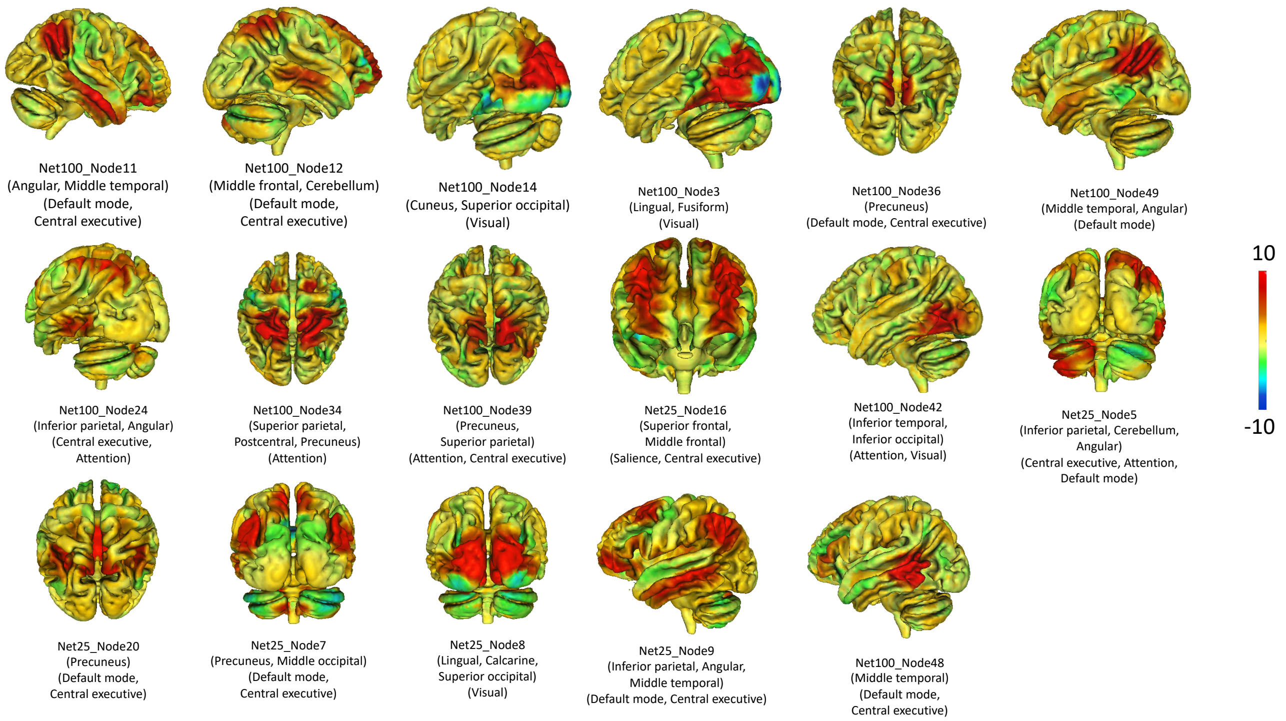


**Supplementary Figure 4:** Location and functional network of the amplitude traits that were associated with the 19q13.32 genomic region.

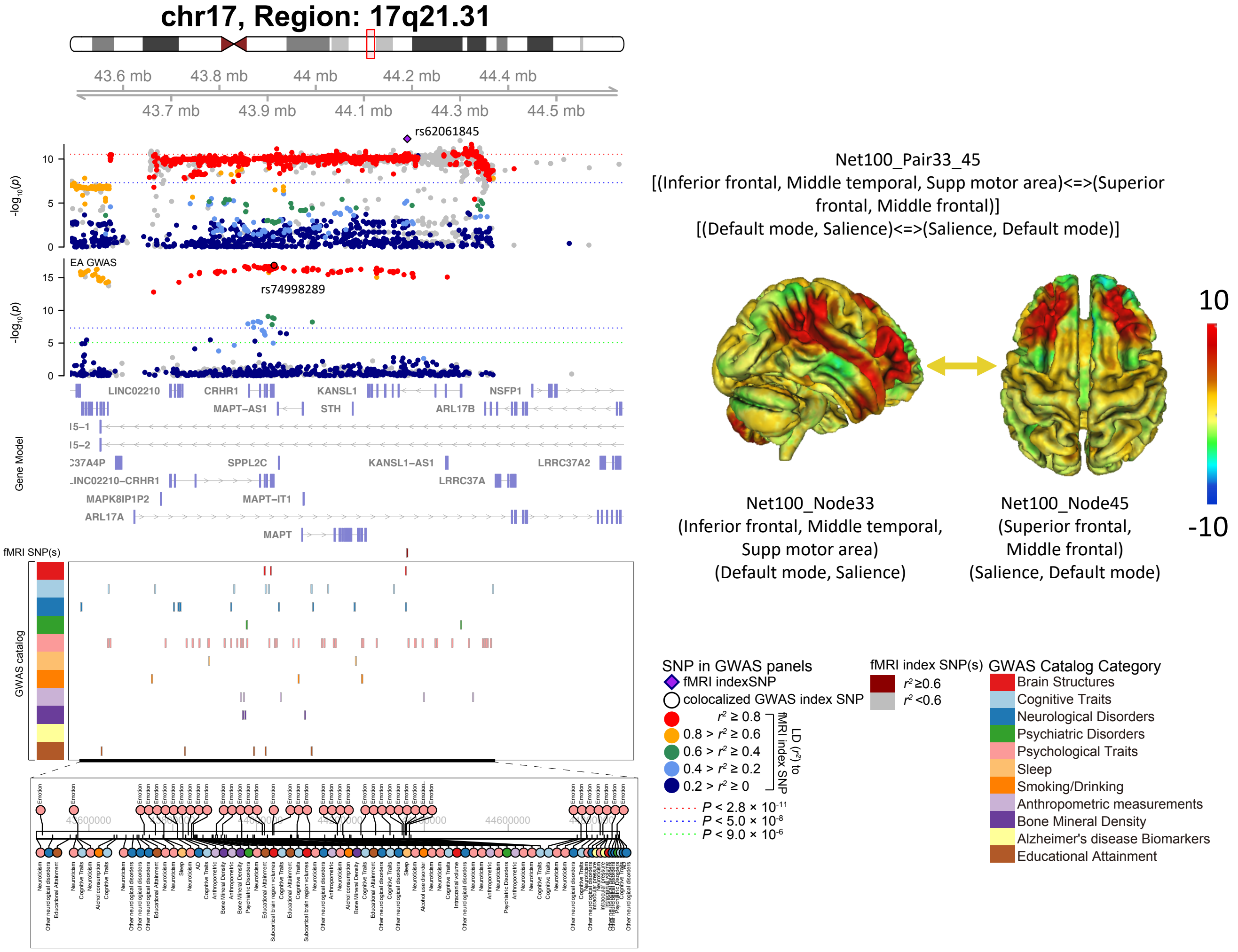


**Supplementary Figure 5:** Local colocalization (LD $r^2 \geq 0.6$) between the selected functional connectivity trait and other brain-related complex traits/disorders in the 17q21.31 genomic region. Location and functional network of the displayed functional connectivity trait are illustrated on the right.

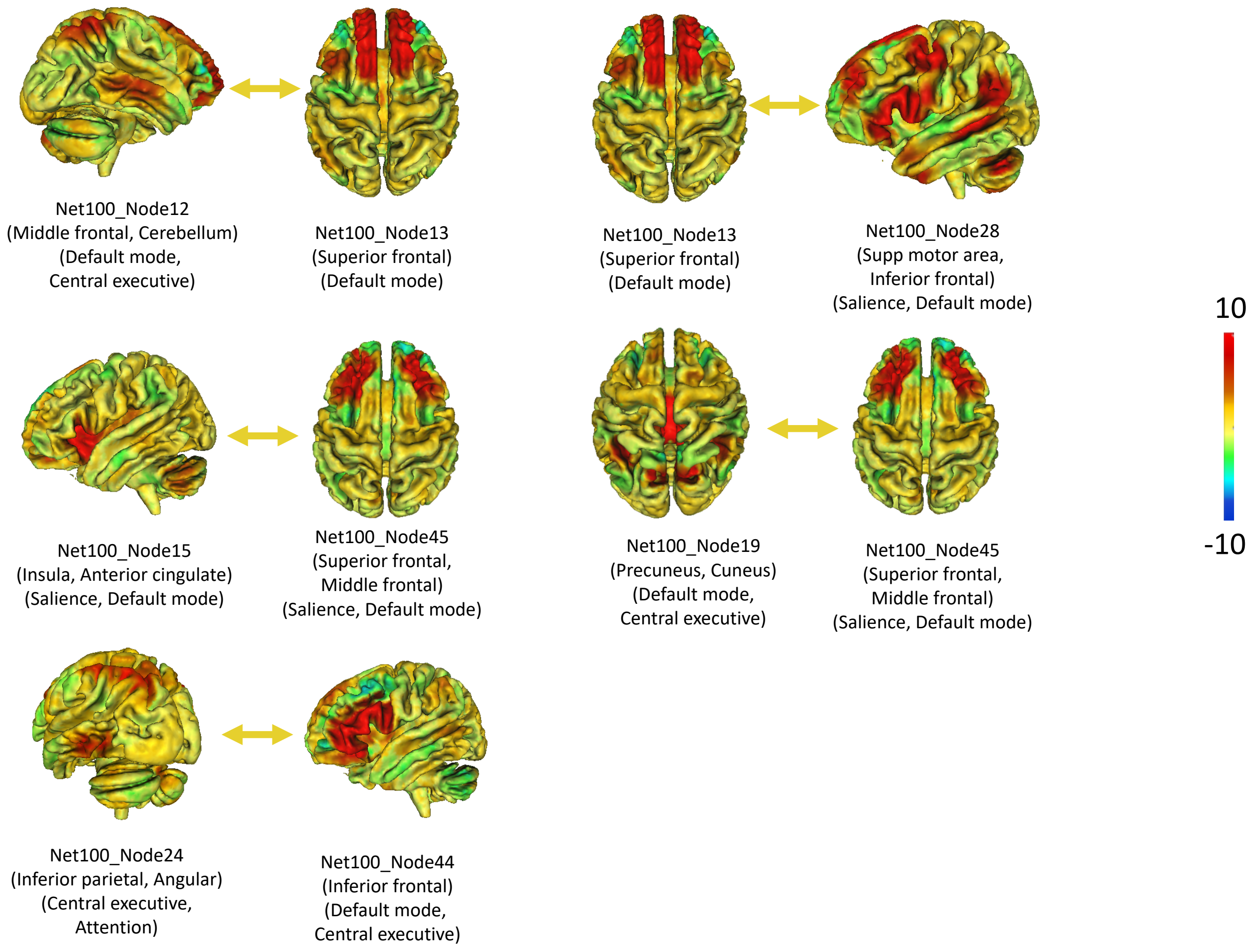


**Supplementary Figure 6:** Location and functional network of the functional connectivity traits that were associated with the 2p16.1 genomic region.

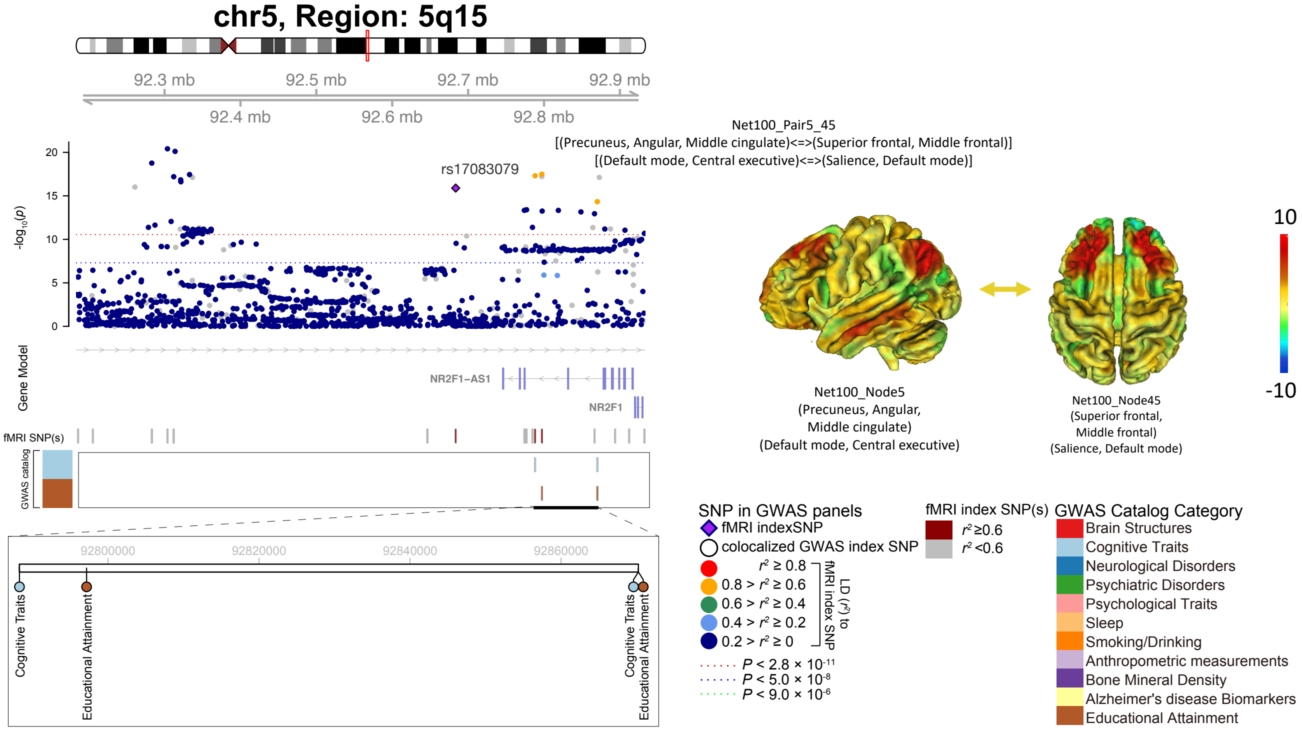


**Supplementary Figure 7:** Local colocalization (LD $r^2 \geq 0.6$) between the selected functional connectivity trait and other brain-related complex traits/disorders in the 5q15 genomic region. Location and functional network of the displayed functional connectivity trait are illustrated on the right.

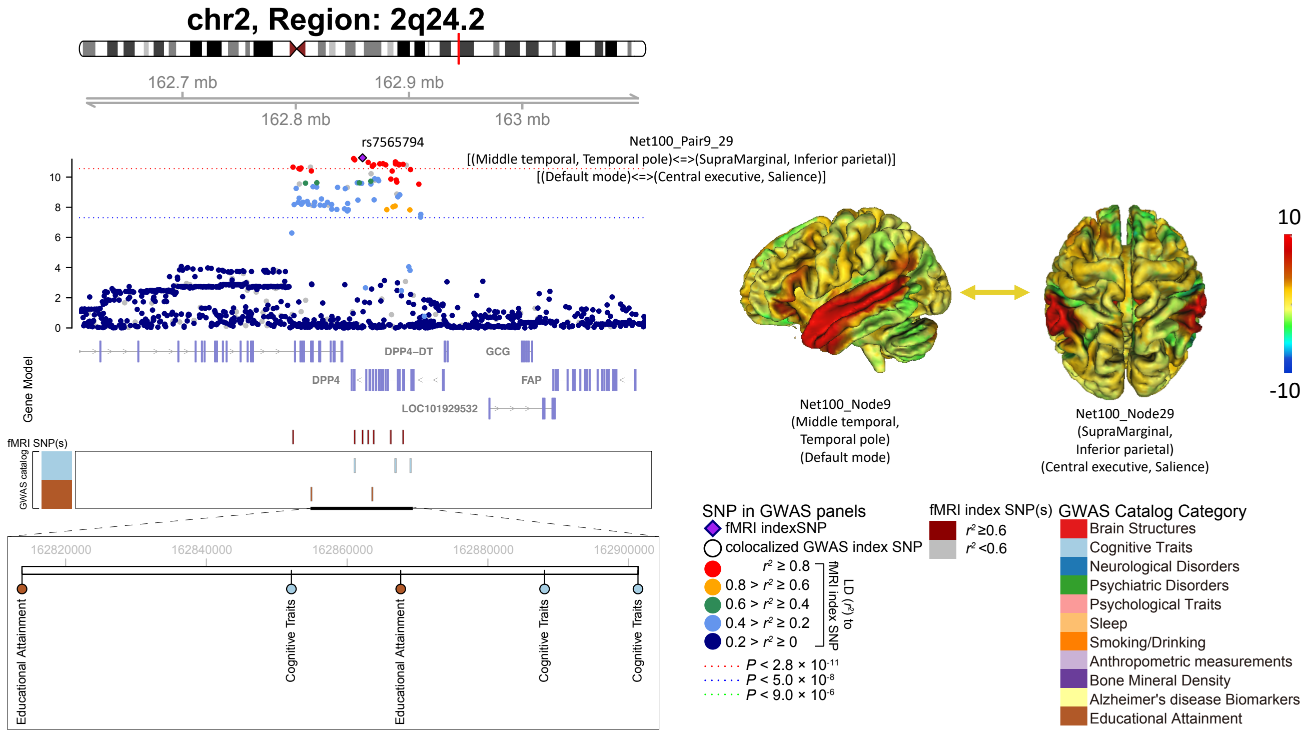


**Supplementary Figure 8:** Local colocalization (LD $r^2 \geq 0.6$) between the selected functional connectivity trait and other brain-related complex traits/disorders in the 2q24.2 genomic region. Location and functional network of the displayed functional connectivity trait are illustrated on the right.

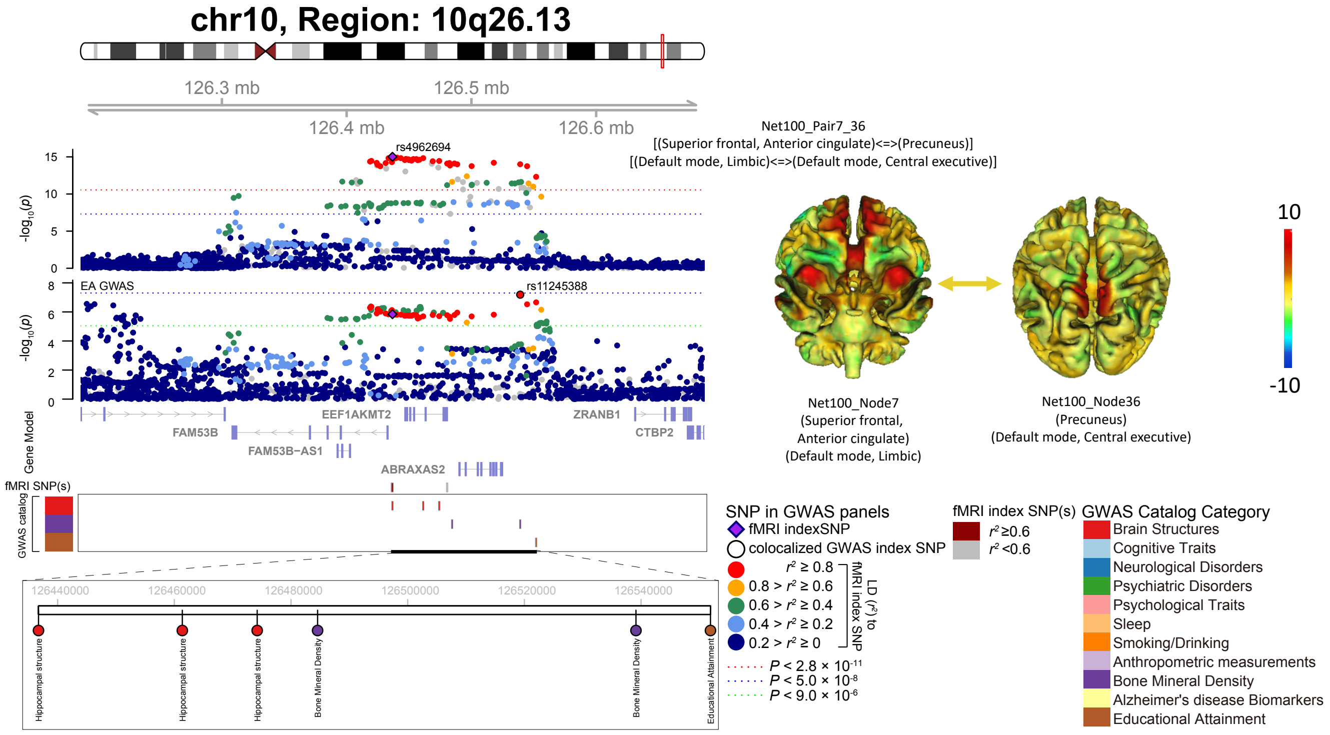


**Supplementary Figure 9:** Local colocalization (LD $r^2 \geq 0.6$) between the selected functional connectivity trait and other brain-related complex traits/disorders in the 10q26.13 genomic region. Location and functional network of the displayed functional connectivity trait are illustrated on the right.

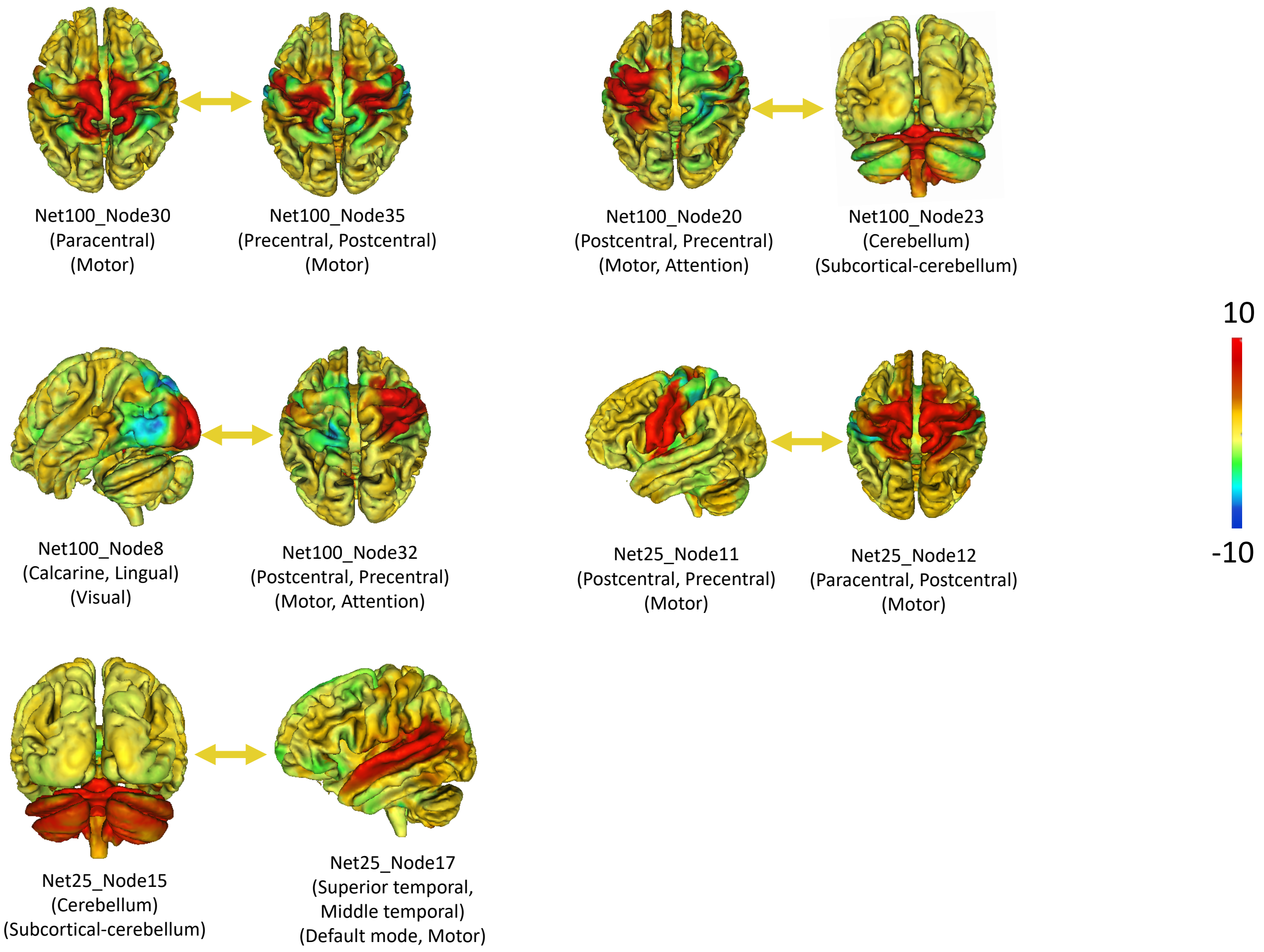


**Supplementary Figure 10:** Location and functional network of the selected functional connectivity traits that were associated with the 2q14.1 genomic region.

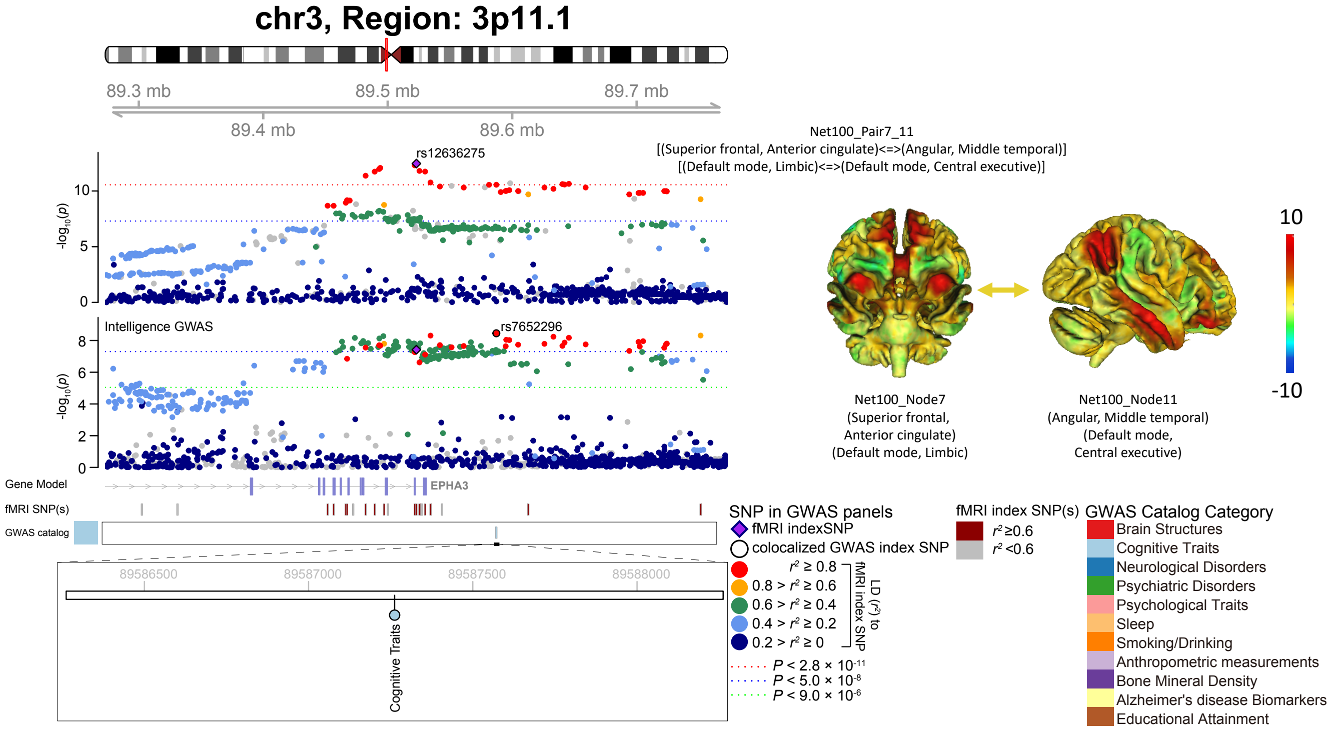


**Supplementary Figure 11:** Local colocalization (LD $r^2 \geq 0.6$) between the selected functional connectivity trait and other brain-related complex traits/disorders in the 3p11.1 genomic region. Location and functional network of the displayed functional connectivity trait are illustrated on the right.

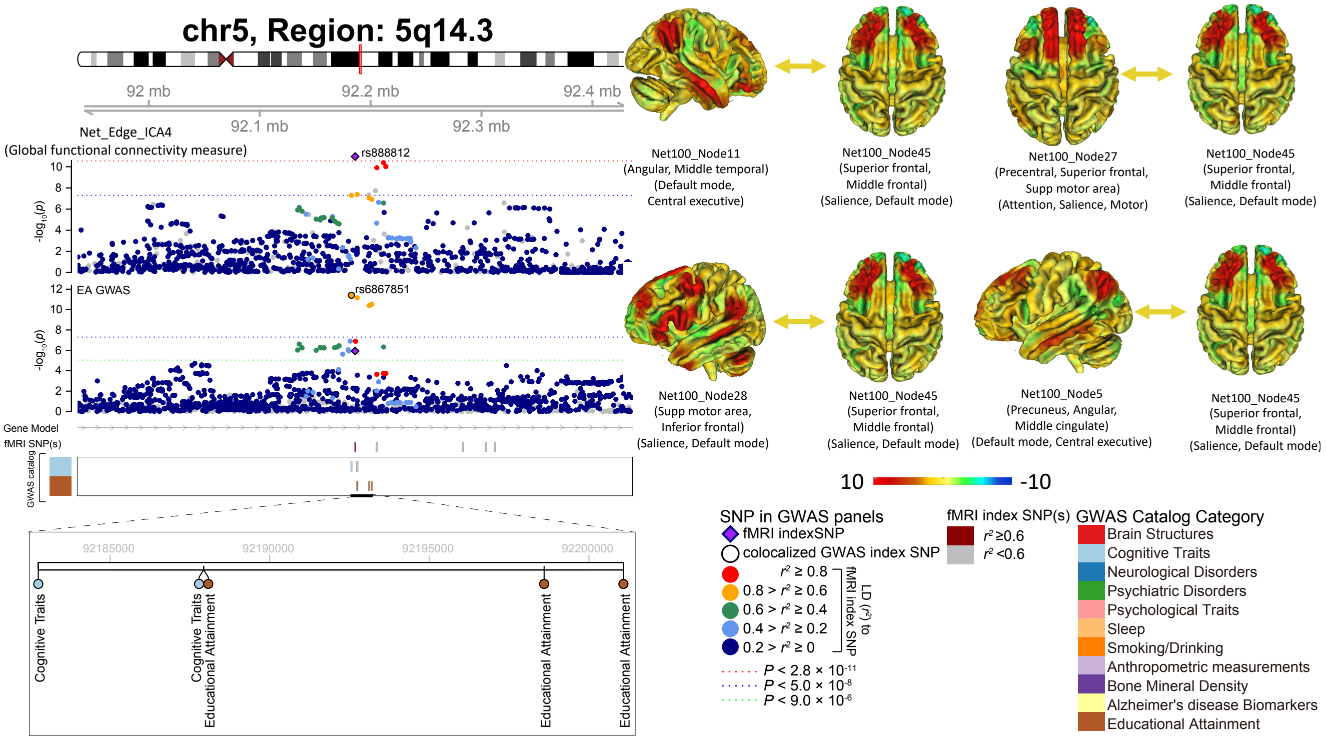


**Supplementary Figure 12:** Local colocalization (LD $r^2 \geq 0.6$) between the selected functional connectivity trait and other brain-related complex traits/disorders in the 5q14.3 genomic region. On the right, we illustrate the location and functional network of the selected pairwise functional connectivity traits that are also associated with 5q14.3.

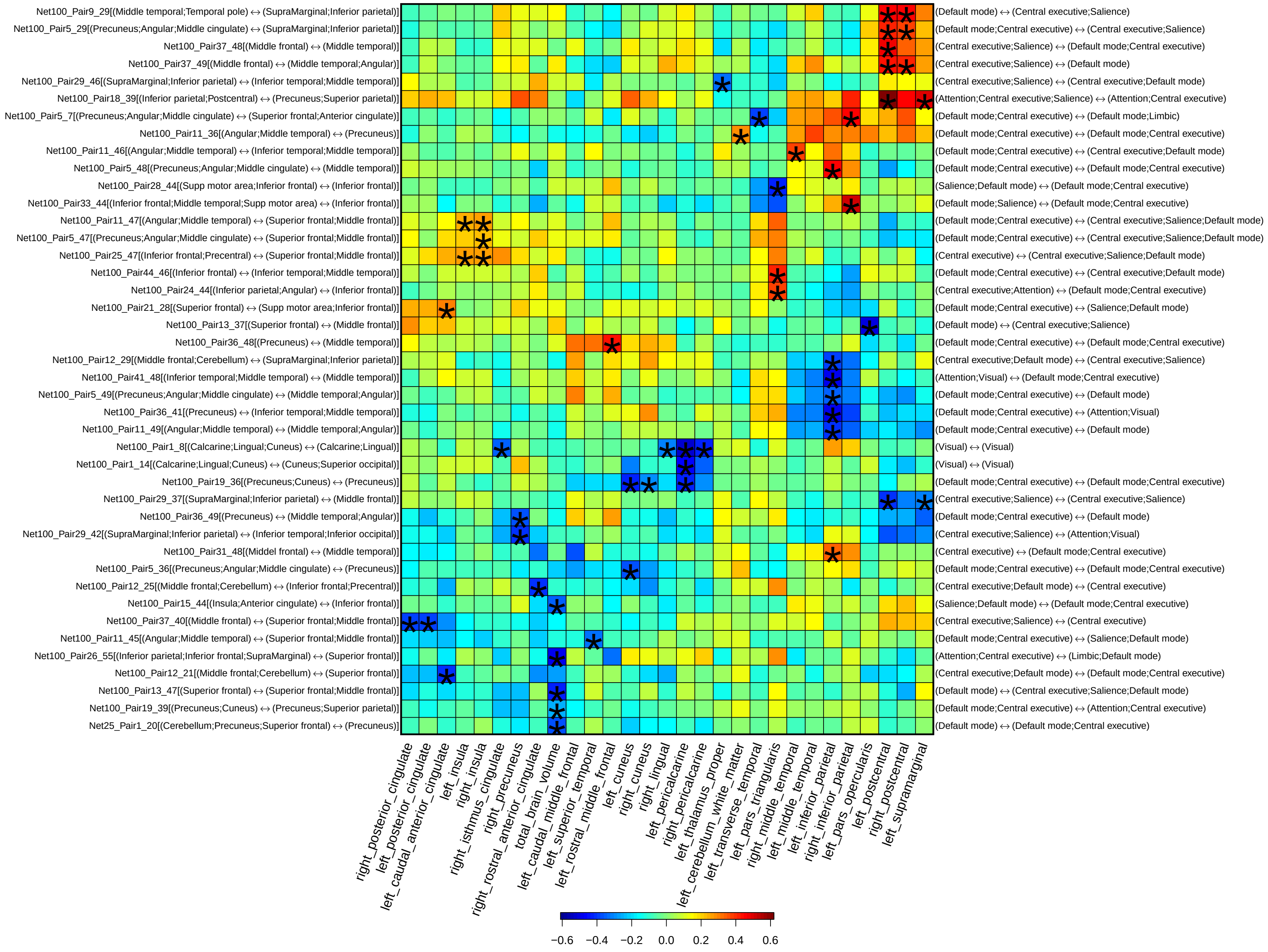

**Supplementary Figure 13:** Selected pairwise genetic correlations between functional connectivity traits and regional brain volumes. The asterisks highlight significant associations after controlling the false discovery rate at 0.05 level (1,777 × 315 tests). The left y axis lists the location of functional connectivity traits, the right y axis shows the associated functional networks, and the x axis provides the name of regional brain volumes.

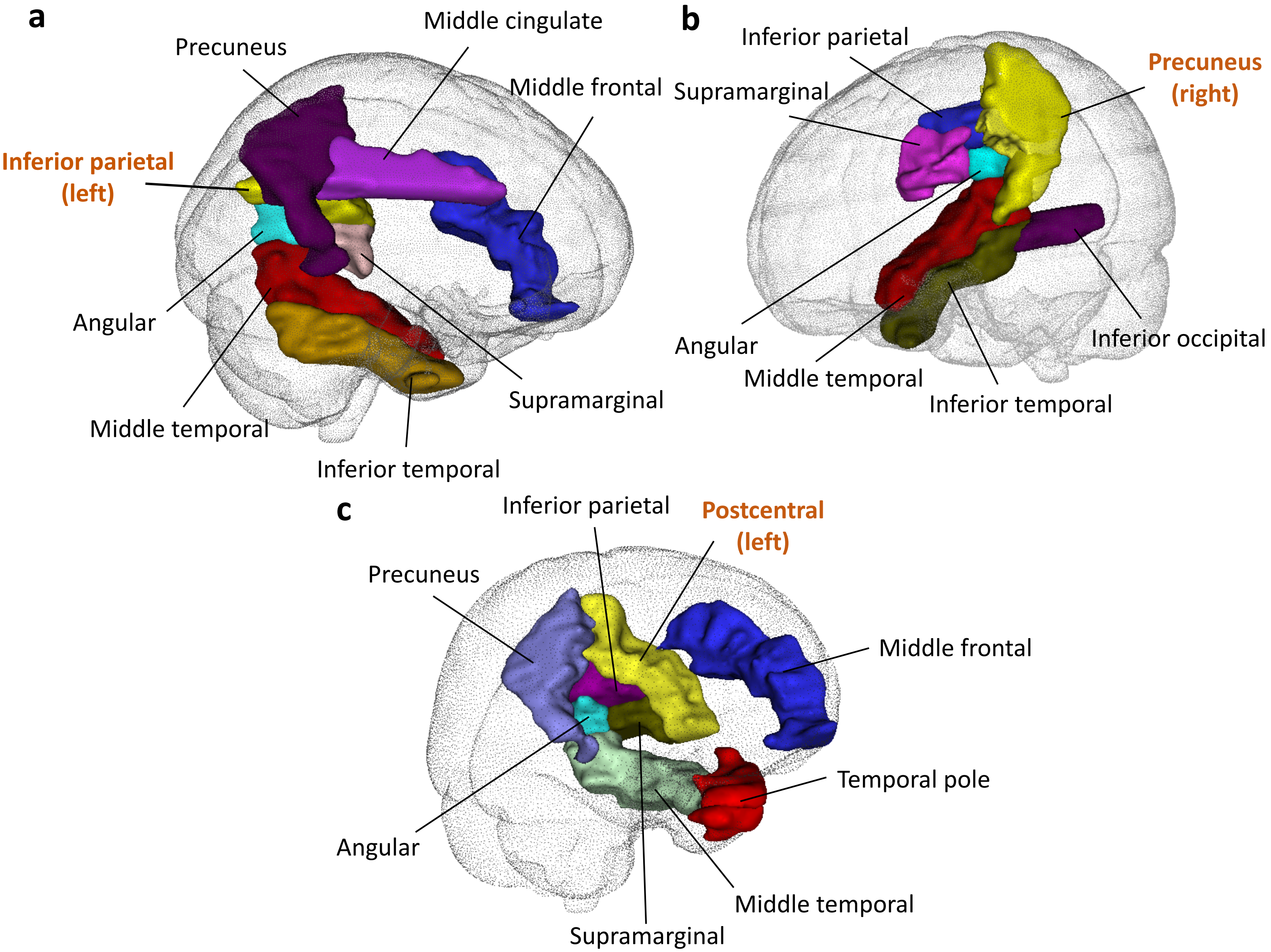


**Supplementary Figure 14:** **a)** Location of the left inferior parietal and its neighboring brain regions whose functional connectivity strengths were genetically correlated with the left inferior parietal volume. **b)** Location of the right precuneus and its neighboring brain regions whose functional connectivity strengths were genetically correlated with the right precuneus volume. **c)** Location of the left postcentral and its neighboring brain regions whose functional connectivity strengths were genetically correlated with the left postcentral volume.

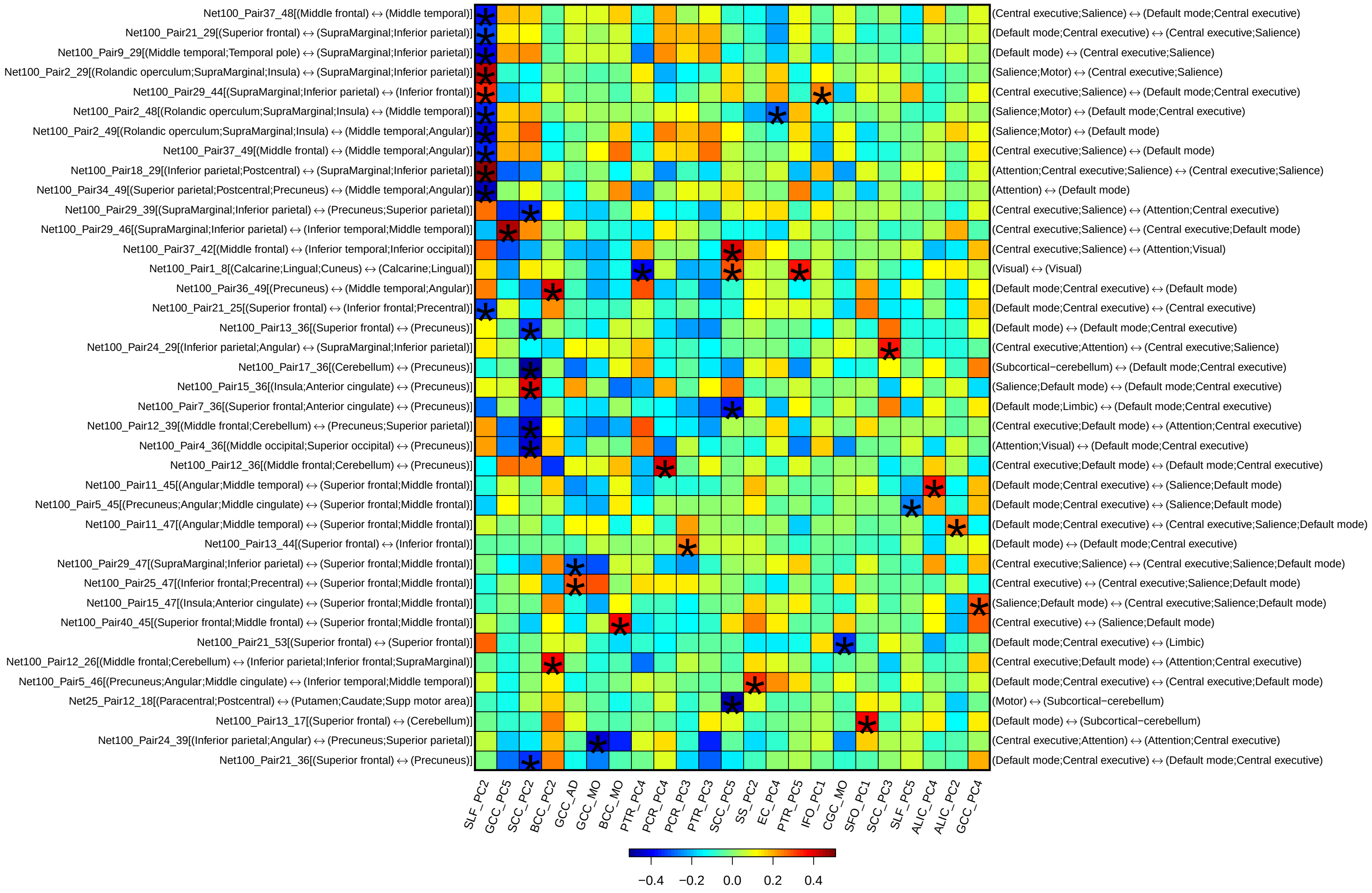


**Supplementary Figure 15:** Selected pairwise genetic correlations between functional connectivity traits and diffusion tensor imaging (DTI) traits of white matter tracts. The asterisks highlight significant associations after controlling the false discovery rate at 0.05 level (1,777 × 315 tests). The left y axis lists the location of functional connectivity traits, the right y axis shows the associated functional networks, and the x axis provides the name of white matter tracts and DTI parameters. SLF, Corticospinal tract; GCC, Genu of corpus callosum; SCC, Splenium of corpus callosum; BCC, Body of corpus callosum; PTR, Posterior thalamic radiation (include optic radiation), PCR, Posterior corona radiata; SS, Sagittal stratum (include inferior longitidinal fasciculus and inferior fronto-occipital fasciculus); EC, External capsule; IFO, Inferior fronto-occipital fasciculus; CGC, Cingulum (cingulate gyrus); SFO, Superior fronto-occipital fasciculus (could be a part of anterior internal capsule); ALIC, Anterior limb of internal capsule. PC, principle component of fractional anisotropy; AD, axial diusivities; MO, mode of anisotropy.

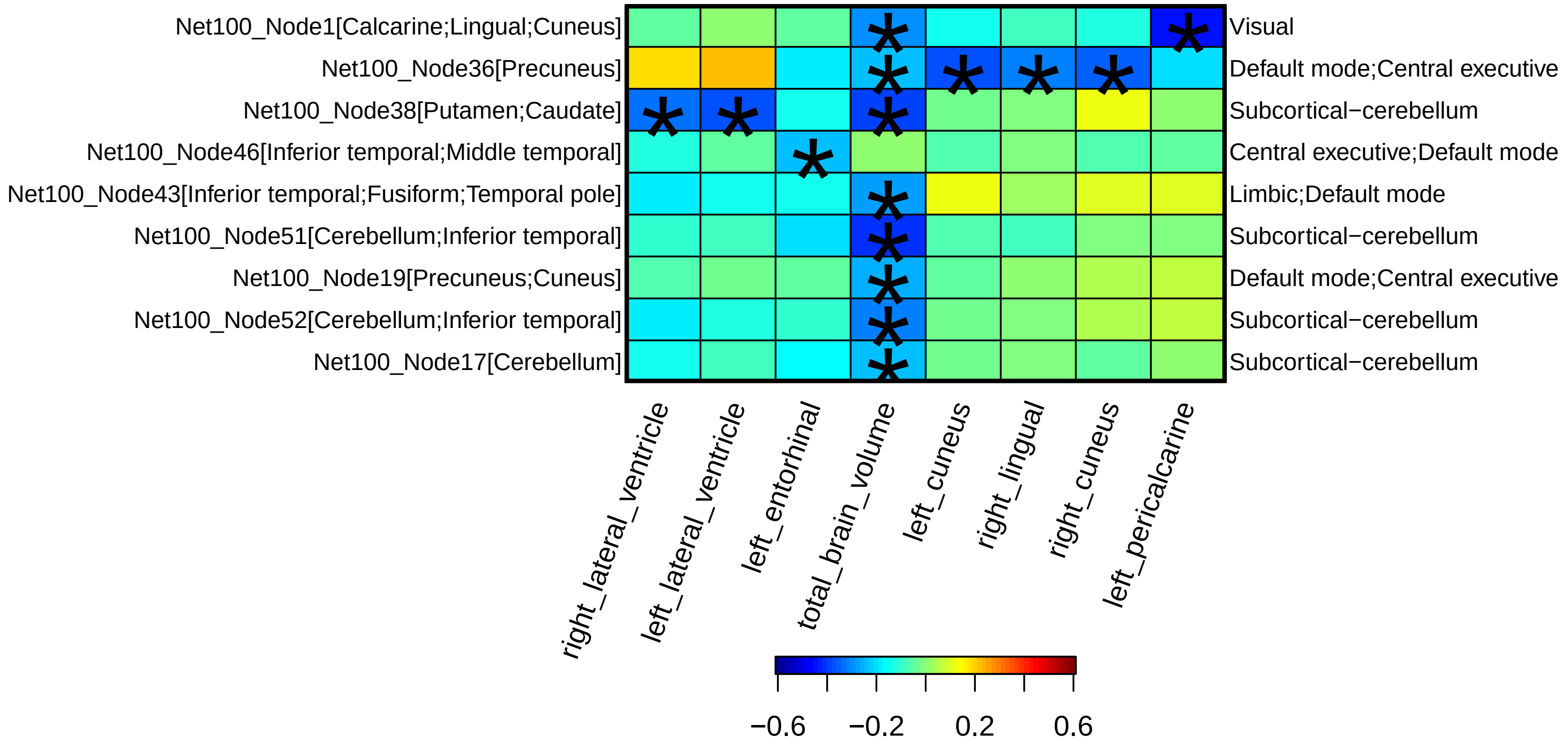


**Supplementary Figure 16:** Selected pairwise genetic correlations between amplitude traits and regional brain volumes. The asterisks highlight significant associations after controlling the false discovery rate at 0.05 level (1,777 × 315 tests). The left y-axis lists the ID and location of amplitude traits, the right y-axis lists the associated functional networks, and the x-axis provides the name of regional brain volumes.

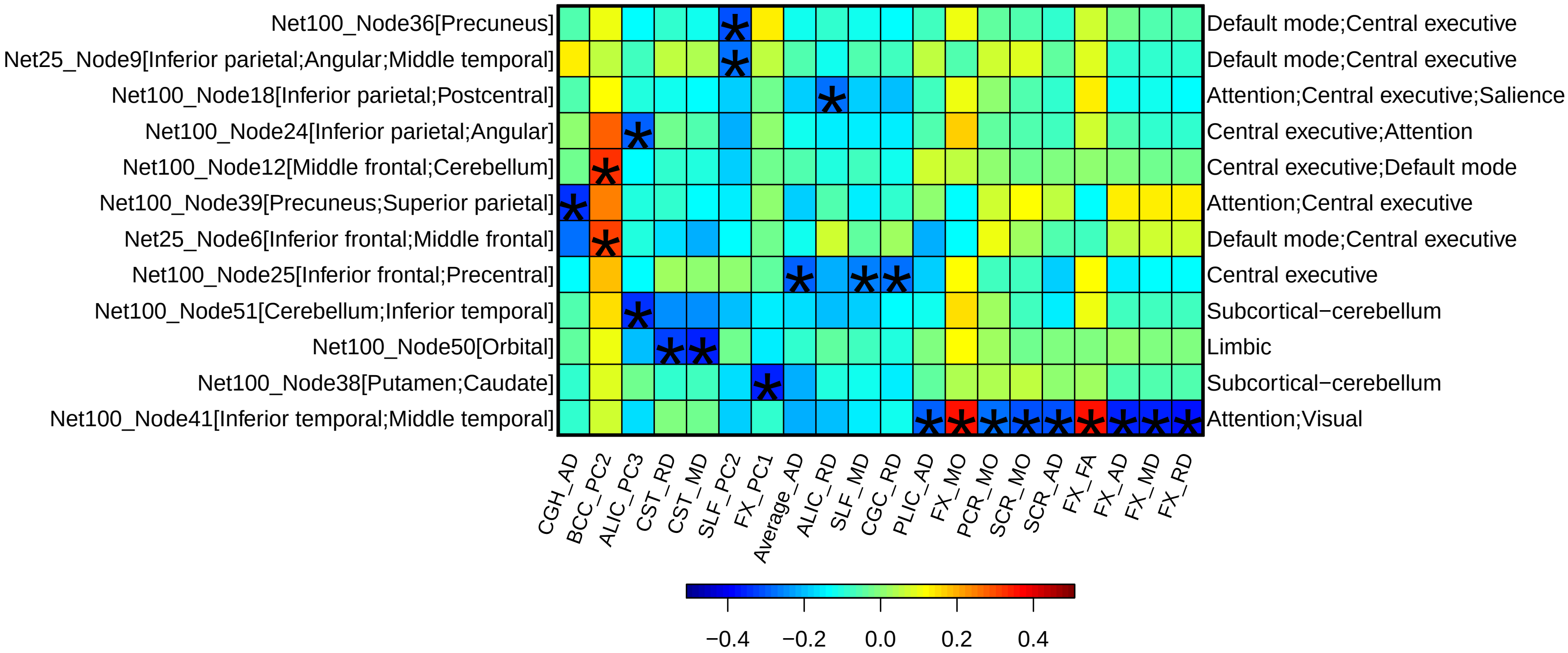


**Supplementary Figure 17:** Selected pairwise genetic correlations between amplitude traits and diffusion tensor imaging (DTI) traits of white matter tracts. The asterisks highlight significant associations after controlling the false discovery rate at 0.05 level (1,777 × 315 tests). The left y axis lists the ID and location of amplitude traits, the right y axis lists the associated functional networks, and the x axis provides the name of white matter tracts and DTI parameters. CGH, Cingulum (hippocampus); BC, Body of corpus callosum; ALIC, Anterior limb of internal capsule; CST, Corticospinal tract; SLF, Corticospinal tract; CGC, Cingulum (cingulate gyrus); PLIC, Posterior limb of internal capsule; FX, Fornix (column and body of fornix); PCR, Posterior corona radiata; SCR, Superior corona radiata. PC, principle component of fractional anisotropy (FA); AD, axial diusivities; MO, mode of anisotropy; RD, radial diusivities; MD, mean diusivities.

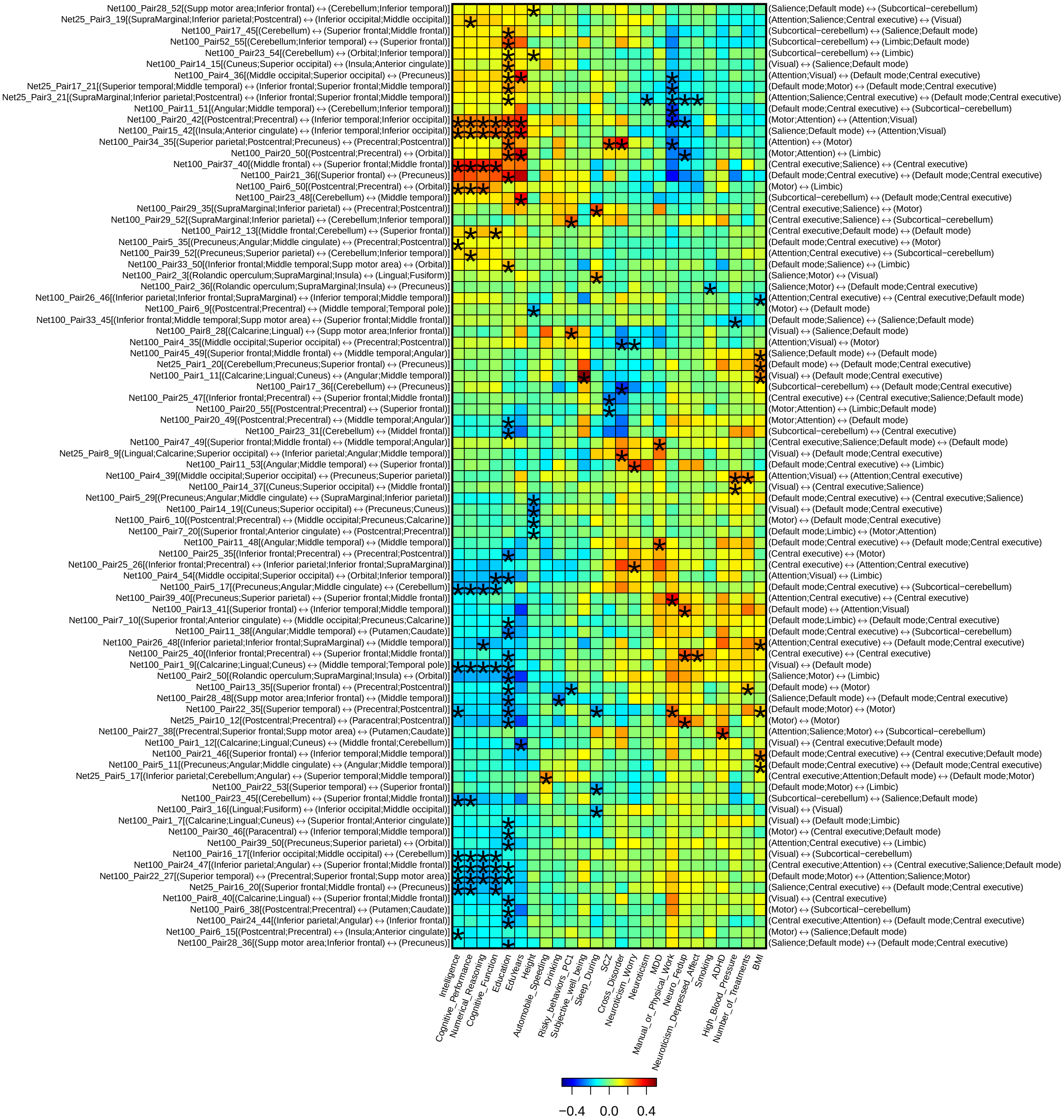


**Supplementary Figure 18:** Selected pairwise genetic correlations between functional connectivity traits and other complex traits. The asterisks highlight significant associations after controlling the false discovery rate at 0.05 level (1,777 × 30 tests). The left y axis lists the ID and location of functional connectivity traits, the right y axis lists the associated functional networks, and the x axis provides the name of other complex traits.

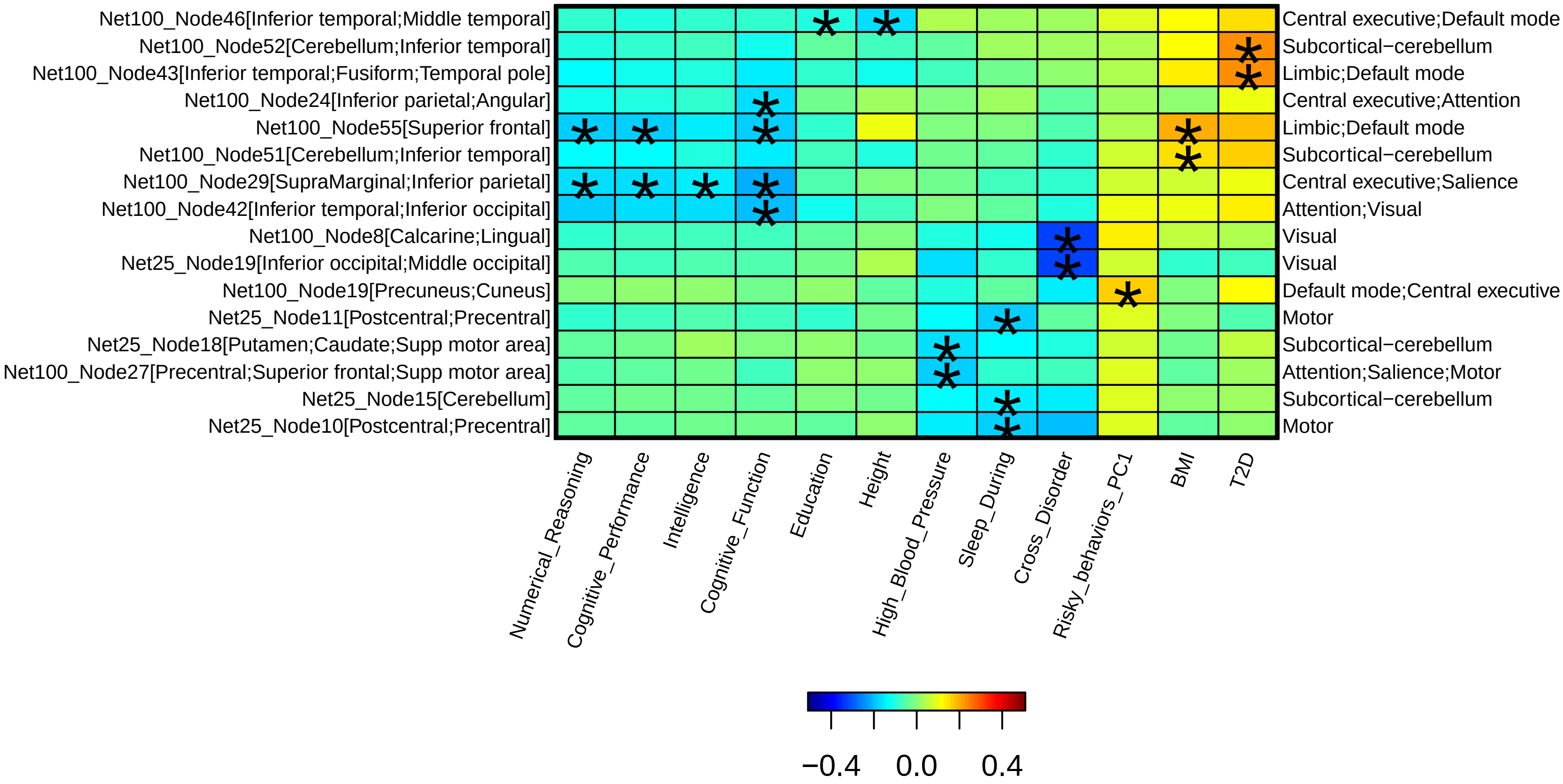


**Supplementary Figure 19:** Selected pairwise genetic correlations between amplitude traits and other complex traits. The asterisks highlight significant associations after controlling the false discovery rate at 0.05 level (1,777 × 30 tests). The left y axis lists the ID and location of amplitude traits, the right y axis lists the associated functional networks, and the x axis provides the name of other complex traits.

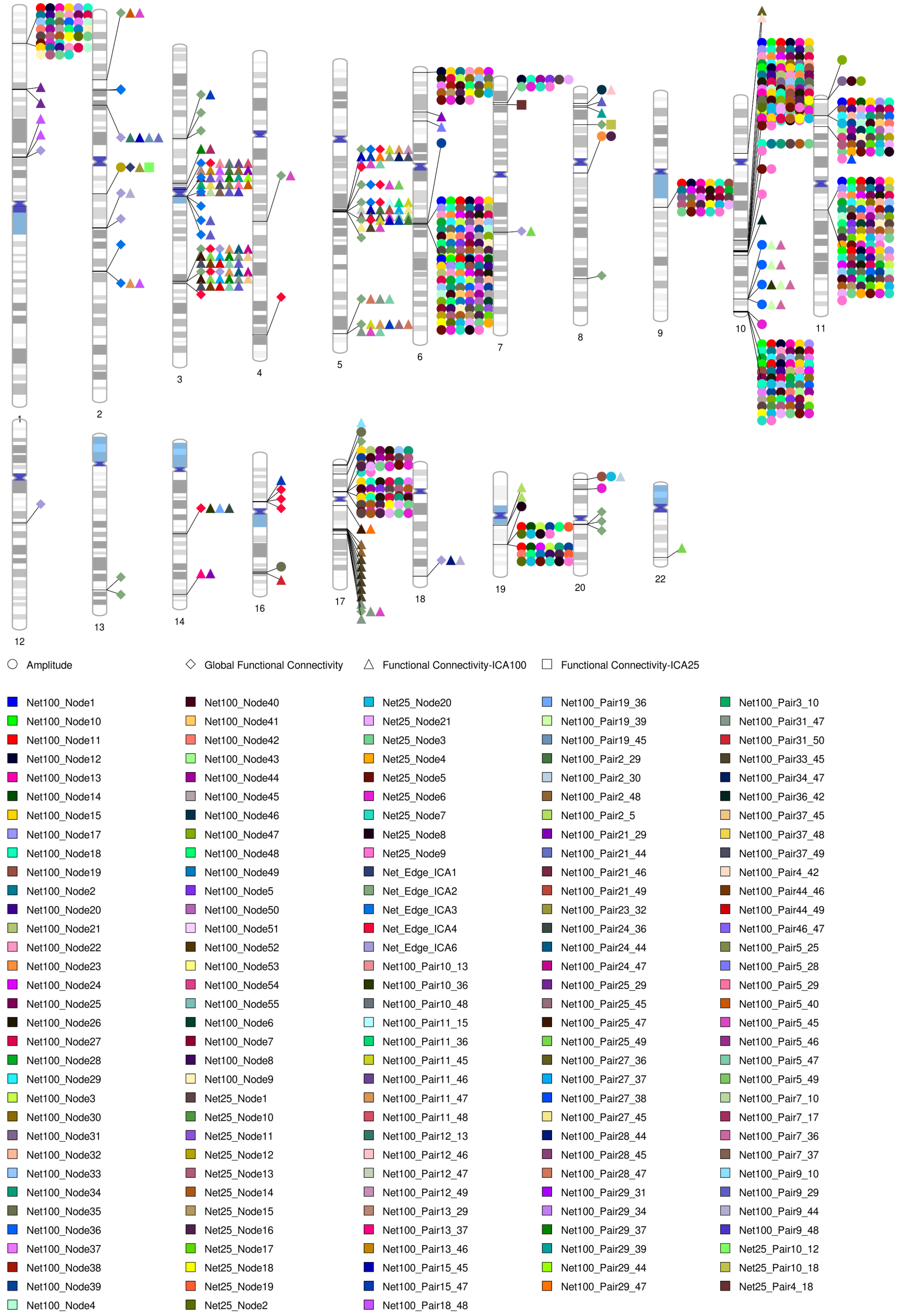


**Supplementary Figure 20:** Location of the genes associated with rsfMRI traits identified in MAGMA gene-level association analysis (n=34,691 subjects).

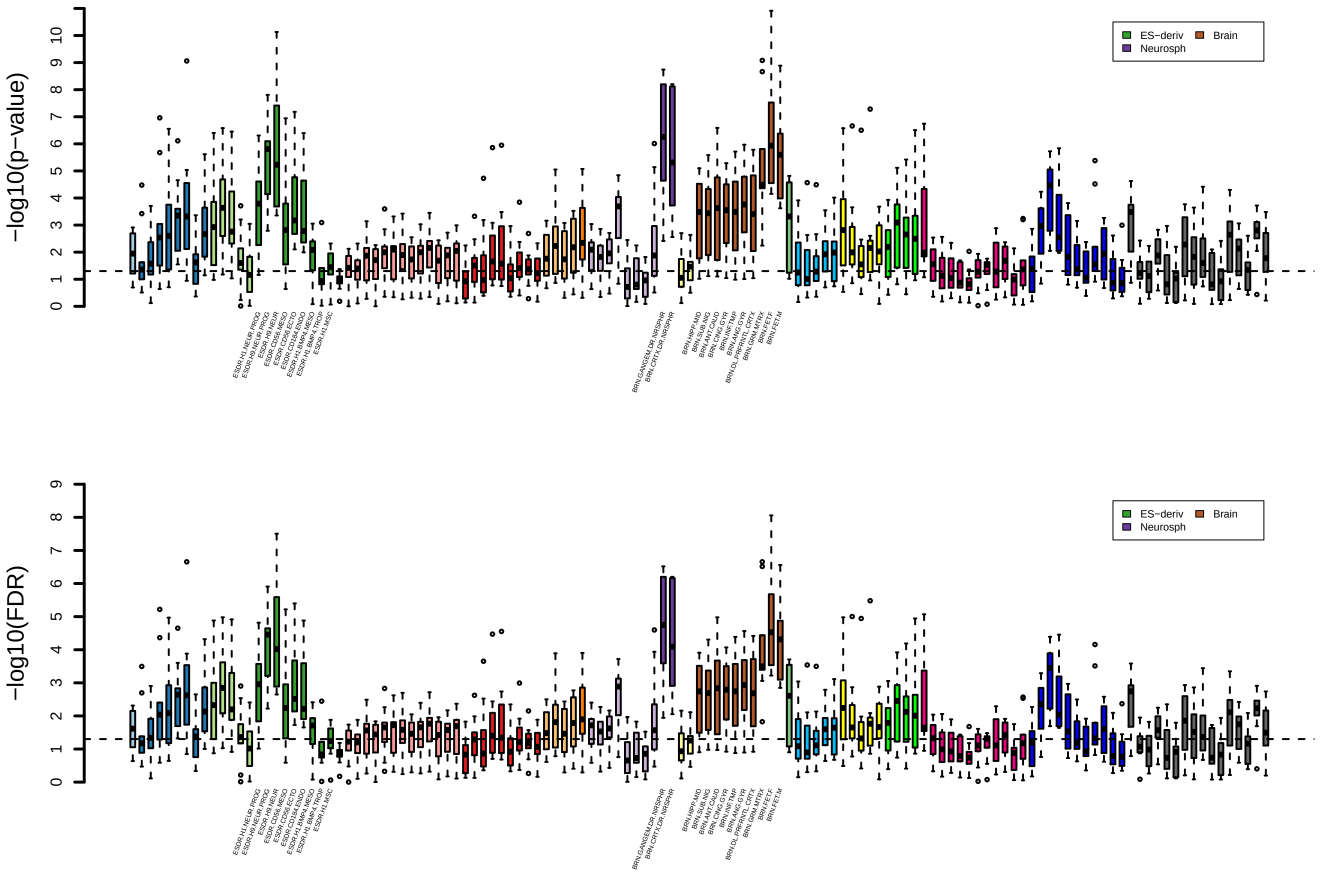


**Supplementary Figure 21:** Partitioned heritability enrichment analysis for tissue type and cell type specific regulatory elements (n = 34,691 subjects). We display the raw p-value (above) and the false discovery rate (FDR) (bottom) in two different panels. The dashed lines indicate the 0.05 significant level. Different colors represent different cell and tissue groups. We label the three most significant tissue and cell types, including fetal brain tissues (Brain), neuronal progenitor cultured cells (ES-deriv), and neurospheres (Neurosph). Enrichment scores and names of all tissue and cell types can be found in Supplementary Table 17.

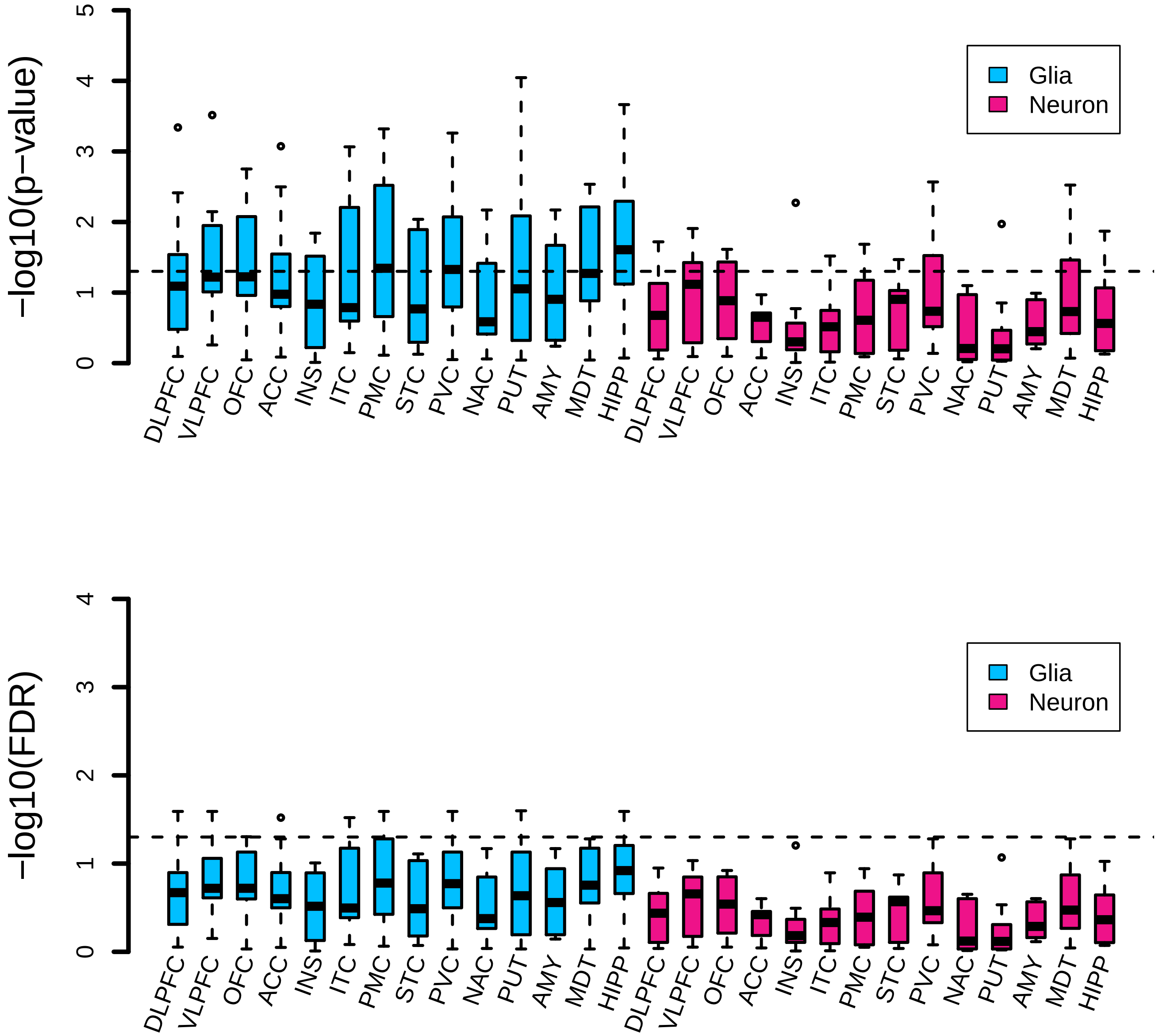


**Supplementary Figure 22:** Heritability enrichment in regulatory elements of two brain cell types (neuron and glia) sampled from 14 brain regions (n = 34,691 subjects). We display the raw p-value (above) and the false discovery rate (FDR) (bottom) in two different panels. The dashed lines indicate the 0.05 significant level. DLFPC, dorsolateral prefrontal cortex; VLPFC, ventrolateral prefrontal cortex; OFC, orbitofrontal cortex; ACC, anterior cingulate cortex; INS, insular cortex; ITC, inferior temporal cortex; STC, superior temporal cortex; STC, primary motor cortex; PVC, Primary visual cortex; AMY, Amygdala; HIPP, Hippocampus; MDT, Mediodorsal thalamus; NAC, Nucleus accumbens; PUT, Putamen.

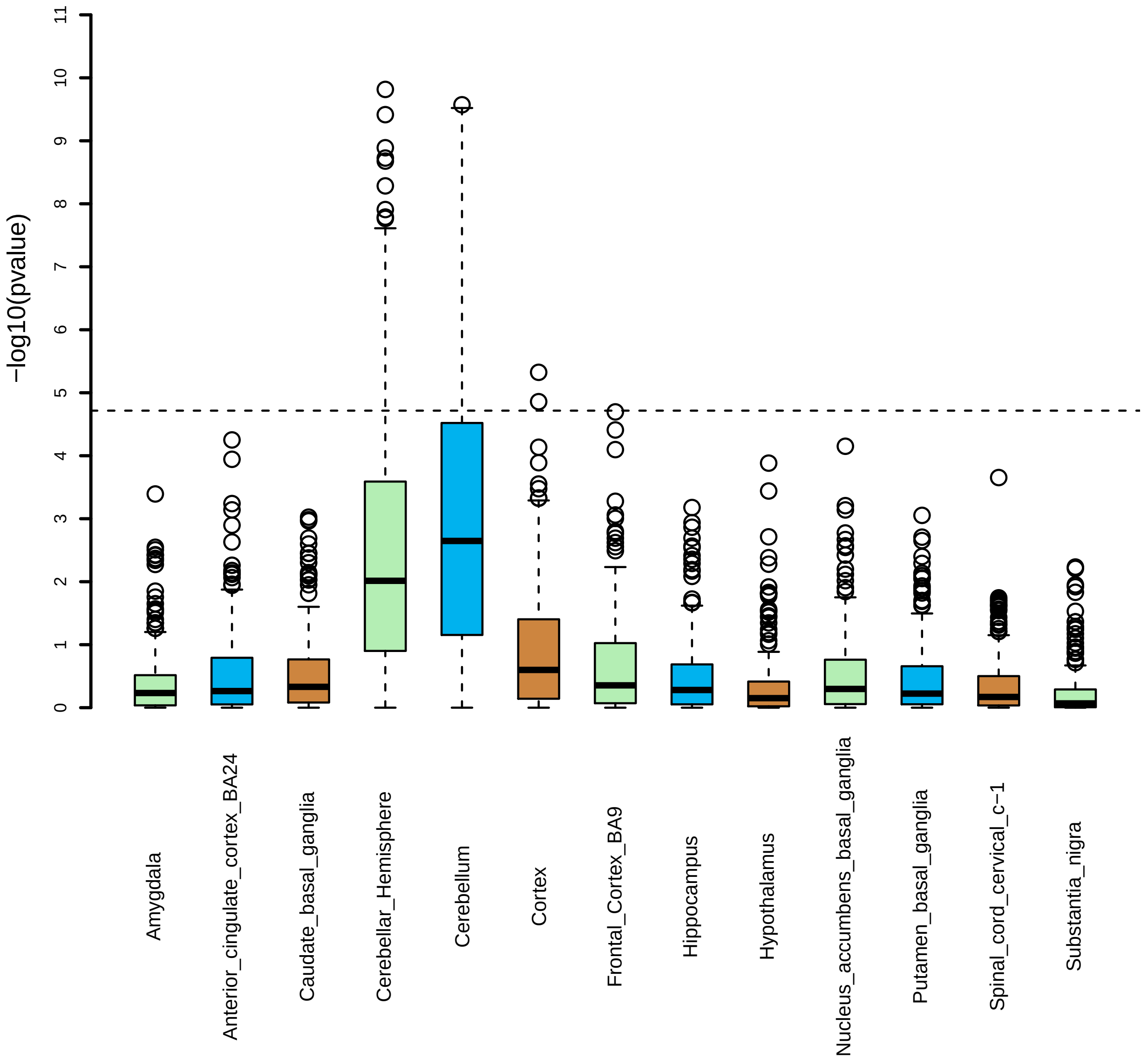


**Supplementary Figure 23:** MAGMA gene property analysis for UKB British discovery GWAS results (n=34,691 subjects) and 13 brain tissues. The 13 brain tissues were from GTEx v8 RNA-seq database. The associations above the horizontal dashed line are significant after Bonferroni correction.

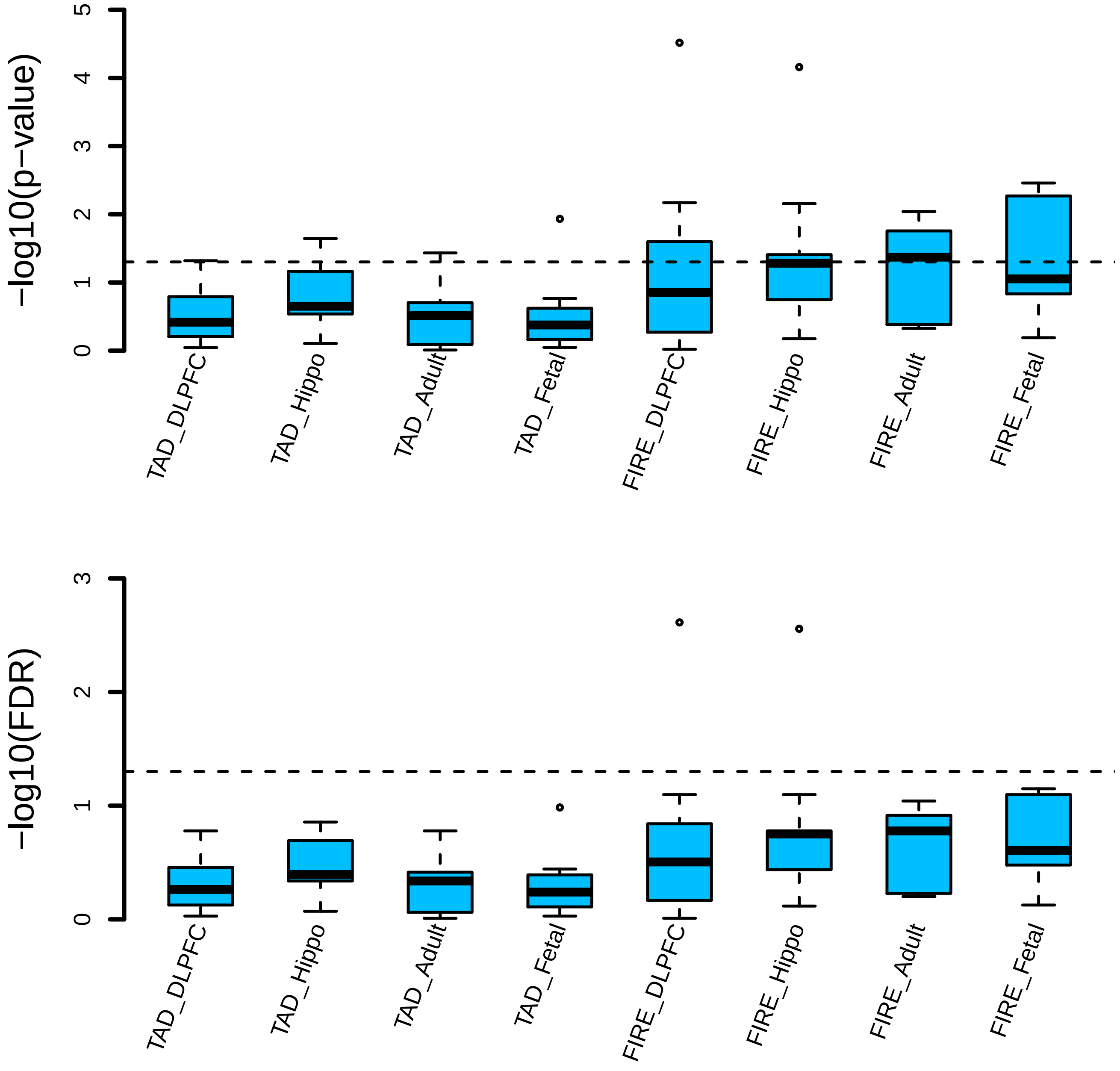


**Supplementary Figure 24:** Heritability enrichment in frequently interacting regions (FIREs) and topologically associating domain (TAD) boundaries of brain tissues (n = 34,691 subjects) We display the raw p-value (above) and the false discovery rate (FDR) (bottom) in two different panels. The dashed lines indicate the 0.05 significant level. DLFPC, dorsolateral prefrontal cortex; Hippo, Hippocampus; Adult, adult brain tissues; Fetal, fetal brain tissues.

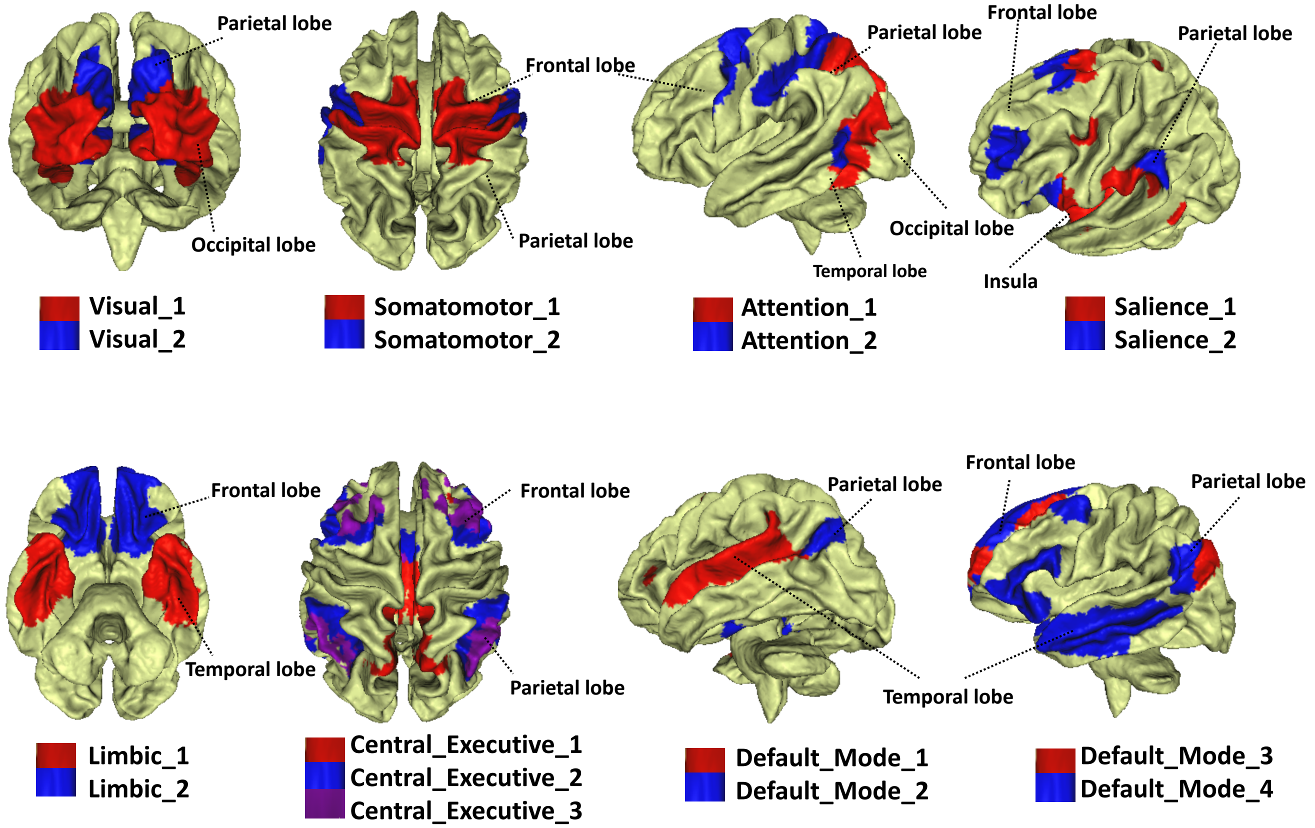


**Supplementary Figure 25:** Annotation of the 17 resting-state networks defined in Yeo et al. (2011).

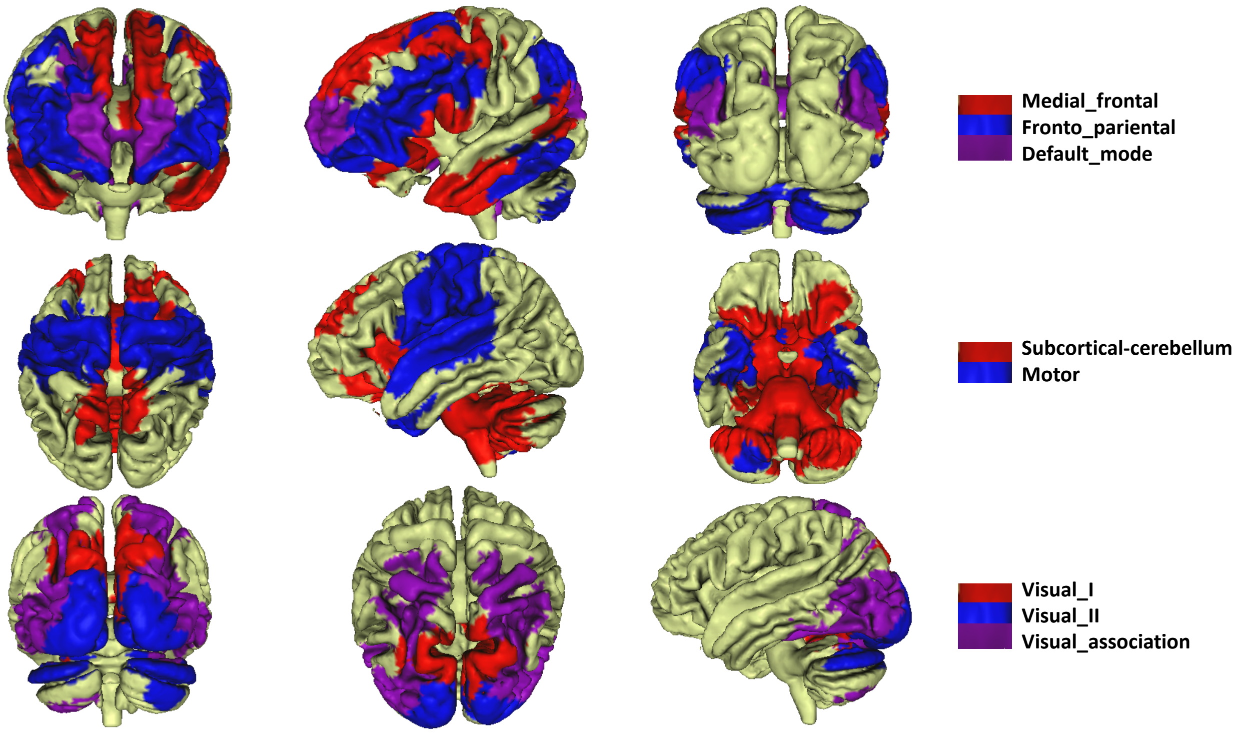


**Supplementary Figure 26:** Annotation of the 8 resting-state networks defined in Finn et al. (2015).
