## Supplementary material for "Common variants contribute to intrinsic human brain functional networks": supp_info

September 10, 2020

#### List of supplementary tables

All supplementary tables can be found in the submitted zip file, here is a list the header of each table.

**Supplementary Table 1:** SNP heritability estimates of 76 amplitude traits and 1701 functional connectivity traits (n=34,691 subjects).

**Supplementary Table 2:** List of independent lead variant-level associations discovered for intrinsic brain activity traits in UKB British discovery GWAS at 2.8e-11 significance level (n=34,691 subjects).

**Supplementary Table 3:** List of significant genetic risk loci discovered for intrinsic brain activity traits in UKB British discovery GWAS at 2.8e-11 significance level (n=34,691 subjects).

**Supplementary Table 4:** Validation p-value of UKB British significant genetic risk loci in European validation datasets (n=6,796 subjects) and non-European validation datasets (n=2,703 subjects).

**Supplementary Table 5:** Incremental R-squared (×100%) and p-value of polygenic risk score for each phenotype constructed by UKB British discovery GWAS (n=34,691 subjects) summary statistics on four European independent datasets.

**Supplementary Table 6:** Incremental R-squared ($\times 100\%$) and p-value of polygenic risk score for each phenotype constructed by UKB British discovery GWAS (n=34,691 subjects) summary statistics on four non-European independent datasets.

**Supplementary Table 7:** Independent significant (p-value $<$ 2.8e-11) variants and their correlated variants for intrinsic brain activity traits that have previously been identified at p-value $<$ 9e-6 in GWAS of any traits listed in the GWAS catalog (version e96_r2019-09-24, www.ebi.ac.uk/gwas/).

**Supplementary Table 8:** Genetic correlation between 1777 intrinsic brain activity traits (n = 34,691 subjects) and 315 brain structural traits, including 100 regional brain volumes and 215 diffusion tensor imaging (DTI) traits of white matter microstructure.

**Supplementary Table 9:** Sources of the publicly available GWAS summary statistics used in this study.

**Supplementary Table 10:** Genetic correlation between 1,777 intrinsic brain activity traits (n = 34,691 subjects) and 30 other complex traits.

**Supplementary Table 11:** List of significant (p-value $<$ 1.5e-9) gene-level associations identified by MAGMA (n=34,691 subjects).

**Supplementary Table 12:** List of mapped genes identified in functional mapping of UKB British discovery GWAS results at 2.8e-11 significance level (n=34,691 subjects).

**Supplementary Table 13:** List of genes associated with intrinsic brain activity traits that have been linked to regional brain volumes, white matter microstructure, and both of them.

**Supplementary Table 14:** Associations of the genes related to all of the three neuroimaging modalities (intrinsic brain activity traits, white matter microstructure, and regional brain volumes) that have previously been report in GWAS of any traits listed in the GWAS catalog (version e98_r2020-02-08, www.ebi.ac.uk/gwas/).

**Supplementary Table 15:** Nervous system drug-target genes that are associated with intrinsic brain activity traits (n=34,691 subjects).

**Supplementary Table 16:** Associations of the genes related to intrinsic brain activity traits that have previously been report in GWAS of any traits listed in the GWAS catalog (version e98_r2020-02-08, www.ebi.ac.uk/gwas/).

**Supplementary Table 17:** LDSC partitioned heritability enrichment analysis in regulatory elements of multiple tissue and cell types (n=34,691 subjects).

**Supplementary Table 18:** LDSC partitioned heritability enrichment analysis in regulatory elements of gross brain cell types (i.e., neurons and glia) and brain cell subtypes, including oligodendrocyte, microglia, and astrocyte, as well as GABAergic and glutamatergic neurons (n=34,691 subjects).

**Supplementary Table 19:** MAGMA gene property analysis for UKB British discovery GWAS results (n=34,691 subjects) and 13 brain tissues. The 13 brain tissues were from GTEx v8 RNA-seq database. Significant tissue groupings after Bonferroni correction are highlighted in bold.

**Supplementary Table 20:** Independent (LD<0.2) significant variants of intrinsic brain activity traits in UKB British discovery GWAS at 2.8e-11 significance level (n=34,691 subjects) that were overlapped with Frequently Interacting REgions (FIREs) and Topologically Associating Domain (TAD) boundaries in brain tissues.

**Supplementary Table 21:** LDSC partitioned heritability enrichment analysis in Frequently Interacting REgions (FIREs) and Topologically Associating Domain (TAD) boundaries in brain tissues (n=34,691 subjects).

**Supplementary Table 22:** List of the genes associated with intrinsic brain activity traits prioritized by gene mapping using 14 recent Hi-C datasets of brain tissue and cell types.

**Supplementary Table 23:** Significant (p-value<1.8e-9) gene sets from MAGMA gene-set analysis for UKB British discovery GWAS results (n=34,691 subjects) after bonferroni correction.

**Supplementary Table 24:** The top ranked regions of the automated anatomical labeling (AAL) atlas in each of the 76 functional brain regions (i.e., nodes) characterized by ICA.

**Supplementary Table 25:** ID, location, and network of 1,701 functional connectivity traits (1,695 pairwise functional connectivity traits and 6 global functional connectivity measures) and 76 amplitude traits.

**Supplementary Table 26:** Sample size, number of genetic variants, and demographic information of each dataset.

### Supplementary Note

#### Genotyping and quality controls

We downloaded the imputed genetic variants data from UKB and HCP data resources, respectively. Genotype imputation was performed locally on the PNC datasets via MACH-Admix (Liu et al., 2013). A full description of the imputation procedures in PNC datasets was detailed supplementary information of Zhao et al. (2019). For the genotype imputation on the ABCD study, we first carried out the following quality control procedures before imputation: 1) exclude subjects with more than 10% missing genotypes; 2) exclude variants with minor allele frequency less than 0.001; 3) exclude variants with missing genotype rate larger than 5%; 4) exclude variants that failed the Hardy-Weinberg test at $1 \times 10^{-9}$ level using only self-identified non-Hispanic white population. We then carried out genotype imputation using the Michigan Imputation Server (`https://imputationserver.sph.umich.edu/`; Das et al. (2016)) and 1000 Genomes Phase 3 (Version 5) reference panel (1000-Genomes-Project-Consortium et al., 2015). Imputed SNPs with a $r^2$-value smaller than 0.3 were removed from the imputation output.

We further performed the following genetic variants data quality controls on each dataset: 1) exclude subjects with more than 10% missing genotypes; 2) exclude variants with minor allele frequency less than 0.01; 3) exclude variants with missing genotype rate larger than 10%; 4) exclude variants that failed the Hardy-Weinberg test at $1 \times 10^{-7}$ level; and 5) remove variants with imputation INFO score less than 0.8.

#### Image acquisition and preprocessing

This work made use of resting-state functional magnetic resonance imaging (rsfMRI) data from four different data resources, which in general had different imaging protocols. Specifically, the image acquisition and preprocessing procedures were detailed in UK Biobank Brain Imaging Documentation (`https://biobank.ctsu.ox.ac.uk/crystal/crystal/docs/brain_mri.pdf`) for the UK Biobank (UKB) study, Casey et al. (2018) for ABCD study, Satterthwaite et al. (2014) for PNC study, and Sotiropoulos et al. (2013) for HCP study. Below we briefly introduce the image acquisition and preprocessing procedures used in each study.

**UKB image acquisition** The rsfMRI data of the UK Biobank were acquired at 490 time points and a duration of 6 minutes, each with a 2.4×2.4×2.4 mm spatial resolution at a dimension of 88×88×64. For image acquisition, the gradient-echo echo-planar imaging (GE-EPI) was adopted with a multiband factor of 8, no iPAT, flip angle 52°, and fat saturation. The echo time (TE) and repetition time (TR) were 39 ms and 735 ms, respectively. As implemented in the CMRR multiband acquisition (Moeller et al., 2010), a separate "single-band reference scan" was also acquired. This had the same geometry (including EPI distortion) as the time series data, but had higher between-tissue contrast to noise, and was used as the reference scan in head motion correction and alignment to other modalities (Alfaro-Almagro et al., 2018).

**UKB image preprocessing** The UKB restfMRI data of about 38,000 subjects (released in 2020) were preprocessed by the UK Biobank brain imaging team (Alfaro-Almagro et al., 2018). The full pipeline can be found in Section 3 of `https://biobank.ctsu.ox.ac.uk/crystal/crystal/docs/brain_mri.pdf`, referred to as UKB preprocessing pipeline in this note. The pipeline generally includes three parts: image cleaning, image registration, and representative time series generation. The source codes have been shared by the UK Biobank team at `https://git.fmrib.ox.ac.uk/falmagro/UK_biobank_pipeline_v_1`.

The image cleaning workflow in the UKB preprocessing pipeline includes the following steps: motion correction using MCFLIRT (Jenkinson et al., 2002); grand-mean intensity normalisation of the entire 4D dataset by a single multiplicative factor; highpass temporal filtering (Gaussian-weighted least-squares straight line fitting, with sigma=50.0s); EPI unwarping; GDC unwarping. Finally, structured artefacts were removed by ICA+FIX processing (i.e., independent component analysis (ICA) followed by FMRIB's ICA-based X-noiseifier (Beckmann and Smith, 2004; Salimi-Khorshidi et al., 2014; Griffanti et al., 2014). FIX was hand-trained on 40 UK Biobank rsfMRI subjects by the UK Biobank team.

The image registration part has the following steps. First, we aligned the GDC unwarped rsfMRI data from the previous step with the high-resolution T1 MRI image. The EPI unwarping in the last step already included an alignment to the T1, though the unwarped data was written out in native (unwarped) fMRI space (and the transform to T1 space written out separately). This T1 alignment was carried out by FLIRT, with a final BBR cost function (Greve and Fischl, 2009). After the fMRI GDC unwarping, a final FLIRT realignment to T1 was applied, which toke into account any shifts resulting from the GDC unwarping. Second, we registered the T1 MR image for each individual to the standard MNI152 2×2×2 mm space. Third, we combined the two image warping together, conducted transformation from the GDC unwarped fMRI space to the MNI standard space, and registered the cleaned fMRI data from the previous step to the MNI standard space by applying the combined image warping. The above three steps were completed in the FMRI expert analysis tool (FEAT) from the software FSL.

Finally, the UKB-derived group-ICA maps including 21 ICA and 55 ICA components were mapped onto the registered cleaned fMRI data to derive the representative time series. These ICA components are publicly available at `http://biobank.ctsu.ox.ac.uk/crystal/refer.cgi?id=9028` and `http://www.fmrib.ox.ac.uk/ukbiobank`. The sets of ICA maps can be considered as "parcellations" of cortical and sub-cortical grey matter, though they lacked some properties often assumed for parcellation. For example, ICA maps were not binary masks but contained a continuous range of values; they can overlap each other; and a given map may include multiple spatially separated peaks/regions. Specifically, these group-ICA maps were obtained by the UK Biobank team using 4,100 subjects through the following procedure: 1) each timeseries dataset was temporally demeaned and had variance normalisation applied according to Beckmann and Smith (2004); 2) group-PCA output was generated by MIGP (MELODIC's Incremental Group-PCA) from all subjects. This comprises the top 1,200 weighted spatial eigenvectors from a group-averaged PCA (a very close approximation to concatenating all subjects' time series and then applying PCA) (Smith et al., 2014); 3) The MIGP output was fed into group-ICA using FSL's MELODIC tool (Hyvärinen, 1999; Beckmann and Smith, 2004), applying spatial-ICA at two different dimensionalities (25 and 100); and 4) 21 out of 25 and 55 out of 100 group-ICA components that were clearly identifiable as artefactual were discarded.

**UKB phenotype generation** The node time series were used to estimate subject-specific network-matrices, which generally included the node amplitude, the Gaussianised full correlation, and partial correlation matrices between node pairs. The correlation-based traits between pairs of brain regions captured the presence of spontaneous co-fluctuations in signal (i.e., the appearance of a connection based on co-activity), while the node amplitude traits reflected the amplitude of spontaneous fluctuation within each region. For each subject, the 21 out of 25 and 55 out of 100 node-timeseries were fed into network modelling. This results in a 21×21 (or 55×55) matrix of connectivity estimates. Network modelling was carried out using the FSLNets toolbox `http://fsl.fmrib.ox.ac.uk/fsl/fslwiki/FSLNets`. The full correlation matrices were derived using fully normalized temporal correlation between every node time series and every other. This was a common and simple approach, but had various practical and interpretational disadvantages, including an inability to differentiate between directly connected nodes and nodes that only connected via an intermediate node (Smith, 2012). Partial temporal correlation IDPs were also calculated between nodes' timeseries, which aimed to estimate direct connection strengths better than the total connection strengths achieved by full correlation. To slightly improve the estimates of partial correlation coefficients, L2 regularization is applied (setting rho=0.5 in the ridge Regression netmats option in FSLNets). Netmat values were Gaussianised from Pearson correlation scores (r-values) into z-statistics, including empirical correction for temporal autocorrelation.

In total 1,695 network-edge functional connectivity traits between many distinct pairs of brain regions were produced in the above step. Next, using the same method of Elliott et al. (2018), we mapped an ICA-based weights matrix to the 1695 IDPs to derive additional 6 ICA features for each individual. The ICA-based weights matrix was online available at `https://www.fmrib.ox.ac.uk/ukbiobank/gwaspaper/`. Specifically, the 6 ICA features were generated by extracting 14 eigenvecters out of the 1695 dimensional IDP matrix using the single value decomposition, followed by the extraction of 6 ICA components out of the 14 eigenvectors using the ICA approach. Robustness of the extracted ICA components were evaluated by the split-half reproducibility approach detailed in Elliott et al. (2018). The resulting six ICA features represented six independent sets (or, more accurately, linear combinations) of the original functional connectivity traits. The selected 21 and 55 nodes as well as matlab code for the above ICA feature generation can be found at `https://www.fmrib.ox.ac.uk/ukbiobank/gwaspaper/`.

By definition of the ICA, the sign of components from ICA are arbitrary. However, the UKB pipeline adopted MELODIC toolbox to calculate the ICA components, which, for convenience of interpretation, applied a simple "skew-related" rule and inverted the signs of the components with negative skewness such that the spatial maps would be dominantly positive.

**Image preprocessing and phenotype generation in other datasets** The ABCD imaging protocol was harmonized for three 3T scanner platforms: Siemens (Prisma VE11B-C), Philips (Achieva dStream, Ingenia), and GE (MR750, DV25-26). This protocol had multi-channel coils that were capable of multiband echo planar imaging (EPI) acquisitions using a standard adult-size coil. The resting-state fMRI data were acquired at 383 time points within a duration of twenty minutes, including eyes open and passive viewing of a cross hair, each with a 2.4×2.4×2.4 mm spatial resolution at a dimension of 90×90×60. The scanning parameters included multiband acceleration factor of 6, flip angle 52°, and the TE and TR being 30 ms and 800 ms, respectively (Casey et al., 2018).

For preprocessing, we downloaded the minimally processed restfMRI dataset, which already went through the following procedure performed by the ABCD team: 1) head motion corrected by registering each frame to the first using AFNI's 3dvolreg; 2) B0 distortions were corrected using the reversing gradient method; 3) displacement field estimated from spin-echo field map scans; 4) applied to gradient-echo images after adjustment for between-scan head motion; 5) corrected for gradient nonlinearity distortions; 5) between scan motion correction across all fMRI scans in imaging event; 6) and registration between T2-weighted, spin-echo B0 calibration scans, and T1-weighted structural images performed using mutual information. Detailes of the above preprocessing steps can be found at the Chapter 15 of the ABCD fix release notes 2.0.1—the NDA 2.0 Resting-State Functional Magnetic Resonance Imaging.

After removal of 8 initial volumes, additional steps were performed on the minimally processed ABCD rsfMRI dataset as follows. First, the ICA+FIX processing was performed to remove structured artefacts to generate the cleaned rsfMRI data. The training subjects for FIX used in the ABCD data were the same as in the UK Biobank data. Similar to the steps of the UK Biobank preprocessing, the cleaned rsfMRI data were then aligned with its corresponding T1 high-resolutional MRI data onto the MNI152 2×2×2 mm space. Next, each time series data were temporally demeaned and had variance normalisation applied. The UKB-derived group-ICA maps including 21 ICA and 55 ICA components were mapped onto the registered ABCD cleaned fMRI data to derive the representative time series on the 76 nodes. Imaging phenotypes including the node amplitude, Gaussianised full-correlation, partial-correlation matrices, as well as the additional 6 ICA features were then generated as we did in the UKB study.

For HCP data, all subjects were scanned on a customized Siemens 3T "Connectome Skyra" scanner housed at Washington University in St. Louis, using a standard 32-channel Siemens receive head coil and a "body" transmission coil designed by Siemens specifically for the smaller space available, as well as the special gradients of the WU-Minn and MGH-UCLA Connectome scanners. The HCP rsfMRI data were acquired in four runs of 14 minutes and 33 seconds each, two runs in one session and two in another session, with eyes open with relaxed fixation on a projected bright cross-hair on a dark background (and presented in a darkened room). Within each session, oblique axial acquisitions alternated between phase encoding in a right-to-left (RL) direction in one run and phase encoding in a left-to-right (LR) direction in the other run. The data were acquired at 1200 time points, each with a 2×2×2 mm isotropic spatial resolution at a dimension of 104×90×72. The gradient-echo echo-planar imaging (GE-EPI) was adopted, with a multiband factor of 8, no iPAT, and flip angle 52°. The echo time and repetition time were 33.1 ms and 720 ms, respectively. The receiver bandwidth was 2290 Hz/Px and the echo spacing was 0.58ms.

The input images of our preprocessing stream were preprocessed HCP rsfMRI images downloaded from the HCP website for the first (RL) and second run (LR) only. Those were both minimally-preprocessed (MPP) and FIX-denoised rsfMRI data, processed by the standard pipeline described in Glasser et al. (2013) and Burgess et al. (2016), and aligned with the corresponding high-resolutional T1 MR images at the MNI152 2×2×2 mm space. Time series data were then temporally demeaned and had variance normalisation applied, and the UKB-derived group-ICA maps including 21 ICA and 55 ICA components were mapped onto the registered ABCD cleaned fMRI data to derive the representative time series on the 76 nodes. Imaging phenotypes including the node amplitude, Gaussianised full-correlation, partial-correlation matrices, as well as the additional 6 ICA features were then generated as we did in the UKB study for the first and second run, respectively. We took the average of the two runs in our downstream analyses.

For PNC dataset, all MRI scans were acquired on a single 3T Siemens TIM Trio whole-body scanner located in the Hospital of the University of Pennsylvania. The system operated under the VB17 revision of the Siemens software. Signal excitation and reception was obtained using a quadrature body coil for transmit and a 32-channel head coil for receive. Gradient performance was 45mT/m, with a maximum slew rate of 200 T/ms. The rsfMRI data were acquired at 124 timepoints within a duration of 6 minutes and 18 seconds, each with a 3×3×3 mm spatial resolution at a dimension of 64×64×46. During the resting-state scan, a fixation cross was displayed as images were acquired. Subjects were instructed to stay awake, keep their eyes open, fixate on the displayed crosshair, and remain still. The scanning parameters included the flip angle 90°, the TE and TR 32 ms and 3000 ms, respectively, and the bandwidth was 2056 HZ per pixel (Satterthwaite et al., 2014). For the preprocessing of PNC dataset, the same pipeline as in the UK Biobank preprocessing, including imaging cleaning, registration, representative time series generation and the IDP generation (except the EPI and GDC unwarping) were applied. The training subjects for FIX used in the PNC data were the same as in the UK Biobank data. Then the imaging phenotypes were generated as we did in the above datasets.

#### Node anatomical location and network classification

The UKB derived group ICA maps include 21 ICA and 55 ICA components (i.e., nodes) on the MNI152 2×2×2mm space after quality controls. The anatomical locations for those ICA maps were detected by the number of voxels with top absolute ICA weights in each region of the AAL atlas (Rolls et al., 2020). Specifically, we focused on the voxels whose nonzero absolute ICA weights were among the top 1%. For each component, we calculated the number of these voxels overlapping with the 170 regions of the AAL atlas. Regions with small overlaps (less than 10 voxels) were removed. For bilateral brain regions, the number of the voxels in the left and right hemispheres were combined. The top ranked regions in each of the 76 ICA node are provided in Supplementary Table 24.

Furthermore, the nodes were classified into 17 brain functional networks defined in Yeo et al. (2011). These networks were shown in Supplementary Figure 25, including two visual, two somatomotor, two attention, two salience, two limibic, three central executive and four default mode networks. Specifically, we first split the 17 functional networks into 34 regions by separating the left and right parts of each network. Then, for the $i$th node, $i = 1, 2, \cdots, 76$, we calculated $Q_{i,j,0.95}$ which was defined as the 95% quantile of the absolute value of its ICA weights within the $j$th region, $j = 1, 2, \cdots, 34$. For each node index $i$, we ranked $Q_{i,j,0.95}$, $j \leq 34$ and picked the regions (as well as networks) with high $Q_{i,j,0.95}$ values and mapped them into the corresponding networks. The 76 nodes were also classified into 8 networks defined by Finn et al. (2015). Those 8 networks consisted of medial frontal, frontal parietal, default mode, subcortical-cerebellum, motor, visual association, and two visual networks (Supplementary Fig. 26). First, the 8 networks were mapped to 268 brain regions defined in Finn et al. (2015). We then calculated $Q_{i,j,0.95}$ for the $i$th node within the $jth$ region, $i = 1, 2, \cdots, 76$, $j = 1, 2, \cdots, 268$. For each node index $i$, we ranked $Q_{i,j,0.95}, j \leq 268$. We picked the regions with high $Q_{i,j,0.95}$ values and mapped them into the corresponding networks.

One of the major differences between the two sets of networks was that Finn et al. (2015) additionally considered the subcortical-cerebellum network. We found that the ICA nodes which had low weights in the 17 networks of Yeo et al. (2011) typically belonged to the subcortical-cerebellum network defined in Finn et al. (2015). Thus, we mainly considered the 17 brain networks from Yeo et al. (2011) and the subcortical-cerebellum network from Finn et al. (2015) in our reported results. The assigned AAL regions and functional networks of the 76 ICA nodes are summarized in Supplementary Table 25.

### More genetic correlation results

#### Amplitude traits and regional brain volumes

We observed significant genetic correlations between amplitude traits and brain volumes (|gc| range = (0.23, 0.44), $P$ range = $(5.2 \times 10^{-12}, 1.4 \times 10^{-5})$, Supplementary Fig. 16). For example, 8 amplitude traits across multiple networks had significant genetic correlations with total brain volume (|gc| range = (0.24, 0.41), $P \leq 1.4 \times 10^{-5}$). It is well known that brain size/volume is phenotypically associated with intrinsic amplitude (Qing and Gong, 2016). Moreover, the amplitude of the putamen and caudate regions in subcortical-cerebellum network was genetically correlated with ventricular volumes. Ventricular volumes are known to be related to subcortical volumes (Okada et al., 2016; Levitt et al., 2002). For the amplitude of precuneus region in default mode and central executive networks, we observed significant genetic correlations with cuneus and lingual volumes. In addition, the amplitude of visual regions (calcarine, lingual, and cuneus) in visual network had significant genetic correlations with the pericalcarine volume. Pericalcarine is involved in the early stage of visual processing (Gomez et al., 2019; Bedny et al., 2012).

#### Amplitude traits and white matter tracts

We detected significant genetic associations between amplitude traits and white matter tracts (|gc| range = (0.27, 0.37), $P$ range = $(1.5 \times 10^{-8}, 1.5 \times 10^{-5})$, Supplementary Fig.. 17). Particularly, our results show that fornix was genetically associated with the amplitude of the middle and inferior temporal regions in the visual and attension networks. Fornix is a critical component of the limbic system and is important in the function of memory (Thomas et al., 2011). For example, the association between the reduced fractional anisotropy in the fornix and performance on visual and spatial memory tests has been found among schizophrenia patients (Fitzsimmons et al., 2009). In addition, we also observed significant genetic correlations between the superior longitudinal fasciculus (SLF) and amplitude in multiple brain regions including the precuneus, inferior parietal, angular, middle temporal, inferior frontal, and precentral. The SLF is involved in a wide variety of brain functions (Klarborg et al., 2012; Hamilton et al., 2008; Rizio and Diaz, 2016; Madhavan et al., 2014; Vestergaard et al., 2011) and is broadly connecting brain regions in temporal, parietal, and frontal lobes (Urger et al., 2015).

#### Functional connectivity traits and schizophrenia

For schizophrenia, we observed significant genetic correlations with connection strengths of central executive, salience, default mode, motor, attention networks, including precentral, postcentral, precuneus, inferior, superior, and middle frontal, and superior parietal regions (|gc| range = (0.18, 0.3), $P$ range = $(3.2 \times 10^{-7}, 1.2 \times 10^{-4})$, Fig. 5a, Supplementary Fig. 18). Hypoconnectivities have been observed over the auditory network (left insula), core network (right superior temporal cortex), default mode network (right medial prefrontal cortex, left precuneus, and anterior cingulate cortices), self-referential network (right superior temporal cortex), and somatomotor network (right precentral gyrus) in schizophrenia patients (Li et al., 2019). The reduced connectivity of postcentral gyrus may play a central role in early-onset schizophrenia (Li et al., 2015). In addition, it has been reported that negative connectivity between language and executive control networks are impaired in schizophrenia patients as well as their first-degree relatives. This decreased connectivity was correlated with performance in language processing (Li et al., 2017).

#### Functional connectivity traits and major depression disorder

For major depression disorder (MDD), significant genetic correlations existed in the middle and superior frontal, angular, and middle temporal regions of the central executive, salience, and default mode networks (|gc| range = (0.26, 0.27), $P$ range = $(1.15 \times 10^{-4}, 1.2 \times 10^{-4})$, Fig. 5a, Supplementary Fig. 18). The temporal and angular gyrus are language-related regions (Ettinger-Veenstra et al., 2016; Dronkers et al., 2011). It has been found that late-onset depression may impair language functions, especially those related to linguistic production (da Silva Novaretti et al., 2011). In addition, altered connectivity strengths in the right angular and the middle temporal have been observed among the treatment-resistant depression and treatment-responsive depression patients (Ma et al., 2012).

#### Functional connectivity traits and subjective well-being

The functional connectivity strength among the calcarine, cuneus, lingual, angular. and middle temporal had strong genetic correlation with subjective well-being (|gc| = 0.48, $P = 2.28 \times 10^{-5}$, Supplementary Fig. 18). Subjective well-being is a scientific term for self-reported happiness and life satisfaction—thinking. The calcarine, cuneus and lingual are the primary visual regions and it has been reported that the primary visual cortex is involved in visual imagery (Kosslyn and Thompson, 2003). The angular is a multimodal convergence hub, which lies at the confluence of brain regions and supports attentional, episodic memory, language and semantic, numerical, and social cognitive processes (Seghier, 2013; Ramanan et al., 2018). Mounting evidence suggests that angular activity scales with subjective ratings of vividness and confidence in recollection, with further evidence pointing to its involvement during construction of detailed and coherent future simulations (Ramanan et al., 2018).

#### Functional connectivity traits and sleep duration

Sleep duration had significant genetic correlations with connection strenghs over auditory (superior temporal), somatosensory (superior parietal, supramarginal), sensory-motor (precentral, postcentral, Rolandic operculum), visual (lingual, fusiform, inferior occipital, middle occipital), insula, and precuneus regions (|gc| range = (0.20, 0.29), $P$ range = $(7.3\times10^{-6}, 1.1\times10^{-4})$, Supplementary Fig.. 18). Horovitz et al. (2008) have demonstrated that blood-oxygen-level-dependent (BOLD) signals increase particularly in visual, motor, and primary auditory cortices when human transits from wakefulness to sleep, which are replicated in other studies (Curtis et al., 2016; Davis et al., 2016; Larson-Prior et al., 2009; Tagliazucchi and Laufs, 2014).

#### Functional connectivity traits and other traits

For high blood pressure, we found genetic correlations with connectivity strengths over the middle occipital, superior occipital, precuneus, superior parietal, cuneus, middle frontal, inferior frontal, superior frontal, middle temporal, and supplementary motor area (|gc| range = (0.19, 0.25), $P$ range = $(2.2 \times 10^{-6}, 8.5 \times 10^{-4})$, Supplementary Fig. 18). It has been reported that participants with hypertension have more activation bilaterally in multiple brain regions, such as the middle occipital, middle temporal, hippocampus, postcentral, insula, and middle frontal (Farcas, 2011).

For risky behavior and automobile speeding, genetic associations mainly existed among motor, central executive, attention, and default mode networks, including the inferior parietal, cerebellum, angular, superior temporal, middle temporal superior frontal, precentral, postcentral, and supramarginal brain regions (|gc| range = (0.20, 0.27), $P \leq 1.5 \times 10^{-4}$), Supplementary Fig. 18) For manual occupation, the genetically correlated brain regions were similar to those associated with cognitive traits and education. Interesting, however, they largely have opposite directions (|gc| range = (0.15, 0.24), $P \leq 1.5 \times 10^{-4}$, Supplementary Fig. 18). Other genetically correlated traits included BMI (|gc| range = (0.2, 0.37), $P \leq 1.5 \times 10^{-4}$) and behavioral factors (drinking and smoking), all of which had been linked to brain functional differences (Kullmann et al., 2012; Shokri-Kojori et al., 2017; Zhou et al., 2017).

#### Amplitude traits and complex traits

For amplitude traits, we detected significant genetic correlations with cognitive traits studied in previous GWAS, including cognitive performance, general cognitive function, intelligence, and numerical reasoning (|gc| range = (0.15, 0.21), $P \leq 1.8 \times 10^{-4}$, Supplementary Fig. 19). We also observed significant genetic correlations between the amplitude of visual area (calcarine, lingual, inferior occipital, middle occipital) with cross disorder (i.e., five major psychiatric disorders) (|gc| range = (0.32, 0.33), $P \leq 9.7 \times 10^{-5}$), and between the regions in motor and suhbcortical-cerebellum networks with sleep (|gc| range = (0.15, 0.18), $P \leq 1.6 \times 10^{-4}$). The association between intrinsic amplitude and cognition, sleep, and brain disorders had been previously reported (Fryer et al., 2015; Meng et al., 2020; Liu et al., 2018).
